# Quantifying asymmetry in non-symmetrical morphologies, with an example from Cetacea

**DOI:** 10.1101/2021.11.17.468940

**Authors:** Ellen. J. Coombs, Ryan. N. Felice

**Affiliations:** Department of Genetics, Evolution, and Environment, University College London, London, UK; Department of Life Sciences, The Natural History Museum, London, UK; Centre for Integrative Anatomy, Department of Cell and Developmental Biology, University College London, London, UK

**Keywords:** asymmetry, geometric morphometrics, landmarks, morphology

## Abstract

1. Three-dimensional measurements of morphology are key to gaining an understanding of a species’ biology and to answering subsequent questions regarding the processes of ecology (or palaeoecology), function, and evolution. However, the collection of morphometric data is often focused on methods designed to produce data on bilaterally symmetric morphologies which may mischaracterise asymmetric structures.

2. Using 3D landmark and curve data on 3D surface meshes of specimens, we present a method for first quantifying the level of asymmetry in a specimen and second, accurately capturing the morphology of asymmetric specimens for further geometric analyses.

3. We provide an example of the process from initial landmark placement, including details on how to place landmarks to quantify the level of asymmetry, and then on how to use this information to accurately capture the morphology of asymmetric morphologies or structures. We use toothed whales (odontocetes) as a case study and include examples of the consequences of mirroring landmarks and curves, a method commonly used in bilaterally symmetrical specimens, on asymmetric specimens.

4. We conclude by presenting a step-by-step method to collecting 3D landmark data on asymmetric specimens. Additionally, we provide code for placing landmarks and curves on asymmetric specimens in a manner designed to both save time and ultimately accurately quantify morphology. This method can be used as a first crucial step in morphometric analyses of any biological specimens by assessing levels of asymmetry and then if required, accurately quantifying this asymmetry. The latter not only saves the researcher time, but also accurately represents the morphology of asymmetric structures.

## Introduction

In recent years there has been a rapid advance in the collection and accumulation of rich morphological data sets using computer tomography (CT) and surface scanning (Davies et al., 2017). High quality data has in turn driven the demand for new methods which accurately and comprehensively capture and represent organismal morphology (Goswami et al., 2019). One such method, geometric morphometrics, often involves the use of 2D or 3D coordinates (landmarks) that are placed on the surface of a specimen or morphology and used to quantify shape independent of isometry, position, and rotation (Bookstein 1991; Zelditch et al. 2004; Lawing and Polly 2010; Adams et al. 2013; Bardua et al., 2019a). Quantifying morphology using these geometric morphometric methods has a long history, and the last few decades in particular has seen an explosion of new advances now used across the biological sciences (Lawing and Polly, 2010; Adams et al., 2013; Bardua et al., 2019a).

In addition to 2D and 3D fixed landmarks, many studies now use semi-landmarks to capture the shape of the regions that fall between landmarks (Gunz et al., 2005; Gunz and Mitteroecker 2013). Semi-landmarks are used to define the outline of structures, such as sutures or ridges, or even entire surfaces, and can provide a significant increase in the shape captured compared to using just landmarks alone (Bookstein, 1991). Curve semi-landmarks (hereafter referred to as ‘curves’ or ‘semi-landmarks’) have been used successfully to quantify a vast array of organismal morphology, including green algae (*Halimeda tuna*: Neustupa and Nemcova, 2018), bird beaks (Cooney et al., 2017), and cranial morphology (Bardua et al., 2019b, Felice et al., 2020). Their use expands the quantification of shape to include the morphology of outlines (e.g., bone margins or veins) and ridges (Cooney et al., 2017; Bardua et al., 2019a). There are numerous reviews which cover the costs and benefits of landmarks alone vs. including semi-landmarks as well as surface semi-landmarks and automated landmarks (see Bardua et al., 2019a).

Coordinate data for structures such as the skull, which is generally bilaterally symmetrical in most species, often comprise landmarks placed on one side of the structure. This is done for several reasons. First, by landmarking only one side and then mirroring those landmarks to the other (symmetrical) side, the user greatly reduces the time required for data capture. Second, landmarking both sides of a symmetrical morphology can produce redundant shape information, but ultimately, using symmetrical data has been shown to improve superimposition (Cardini, 2016a; 2016b). To accommodate these issues, Bardua et al., (2019a) recommend imputation of the missing side through mirroring of the existing landmarks along a midline plane, then removing these mirrored landmarks after Procrustes superimposition (translation to a common origin, scaling to unit centroid size, and rotation to leave only shape data; Mitteroecker and Gunz, 2009) to reduce data dimensionality and redundant information. While this method has proven to work well in capturing the shape of bilaterally symmetrical morphologies (e.g., Adams et al., 2013; Dumont et al., 2016; Felice and Goswami, 2018; Bardua 2019b; Watanabe et al., 2019) into which most aspects of vertebrate anatomy fall, the same cannot be said for rarer asymmetrical morphologies (Fig. 1).

**Fig. 1.**
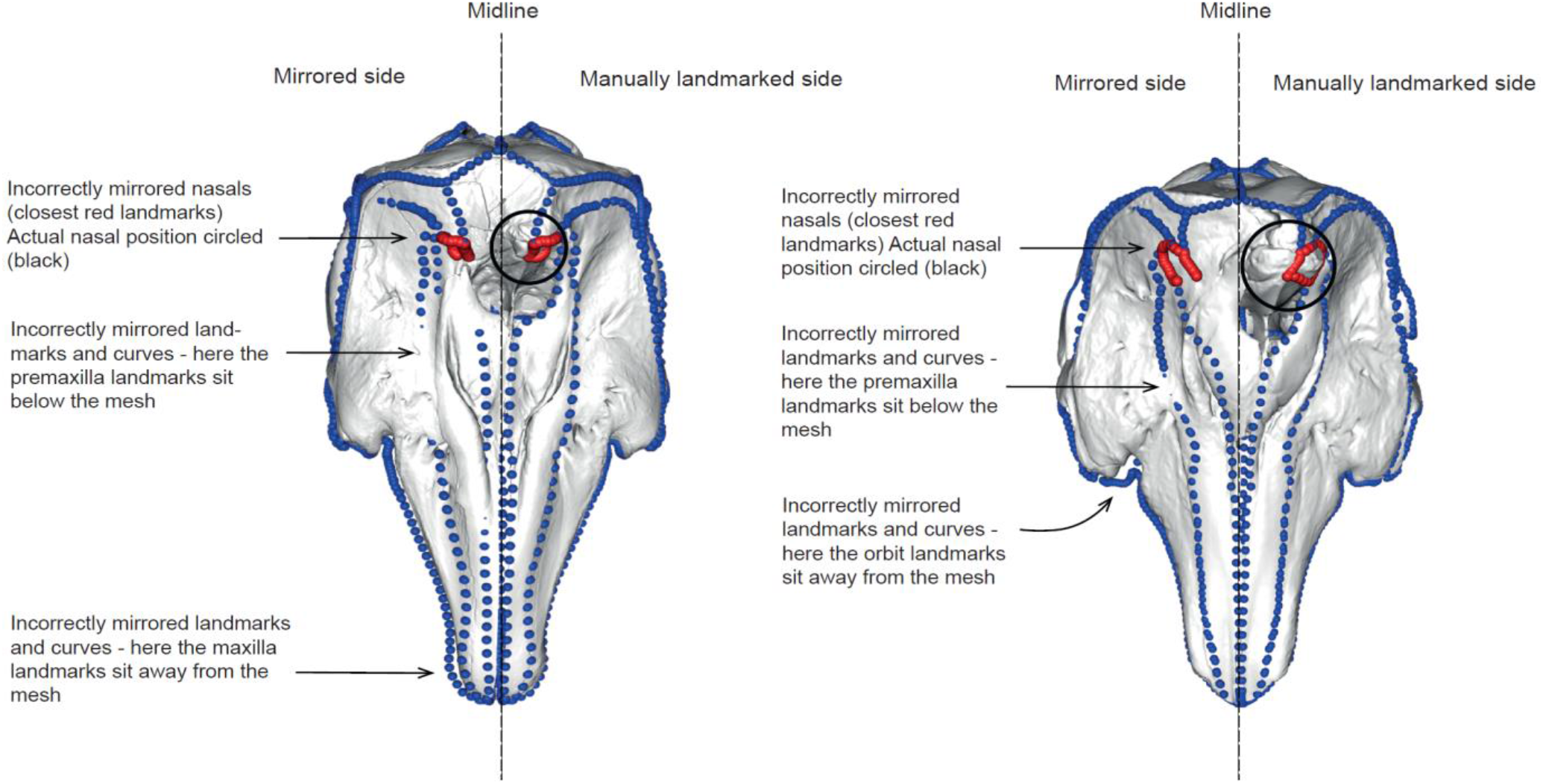
Computer mirrored landmarks can misrepresent true morphology in asymmetric specimens. Landmarks and semi-landmarks are shown in blue on the skulls of two odontocetes. Nasal landmarks are shown in red to illustrate the inaccurate capture of the asymmetric morphology. Nasals and their actual positions are circled in black – ideally all red landmarks should sit within the black circle. Specimens shown are *Delphinapterus leucas* (USNM 305071, left) and *Monodon monoceros* (USNM 267959, right).

### Asymmetrical morphologies

Symmetry in external morphology is the general rule among plants and animals, making the cases of directional asymmetry particularly interesting. However, it is now well known that some organisms do have a naturally occurring asymmetrical morphology. Directional asymmetry (DA), a type of asymmetry that occurs in a consistent direction between a pair of morphological structures, is often related to function or developmental morphology. This is typically expressed as size differences in bilaterally paired structures (e.g., limbs, muscles). This differs from fluctuating asymmetry (FA) which is often used as a measure of stress in populations, of individual quality or of developmental instability (Graham et al., 1993; Klingenberg, 2003). FA is often minute and requires the capturing of measurement error on a different scale to the asymmetry covered in this study.

Directional asymmetry related to function is found across plants and animals with examples including the shells of turtles (yellow-bellied sliders (*Trachemys scripta scripta*); Parés-Casanova, 2020) and the appendicular skeleton of some cetaceans. In some cetaceans, the humerus and ulna is significantly larger on the right (dextral) side in the harbour porpoise (*Phocoena phocoena*) and the white-beaked dolphin (*Lagenorhynchus albirostris*) with a larger dextral muscle mass and higher mechanical stress indicating lateralized behaviours (Galatius, 2005; 2006). Handedness and associated directional asymmetry of limbs is also detected in humans (*Homo sapiens*; Auerbach and Ruff, 2006) and rhesus macaques (*Macaca mulatta*; Falk et al., 1998), and in the pectoral appendages or walruses (*Odobenus rosmarus*; Levermann et al., 2003). In some taxa, such as the fox (*Vulpes vulpes*), directional asymmetry in the limbs, skull, and pelvis is not associated with preferential use but instead with differential biases in growth (Kharlamova et al., 2010).

In contrast to these size-based examples of asymmetry, some taxa exhibit directional asymmetry in the morphology or position of paired structures. In owls (Strigiformes) (Krings et al., 2020), pronounced bilateral asymmetry in the external ears is related to directional hearing, or sound localization ability (Payne I97I; Norberg, 1977; Norberg 2002; Krings et al., 2020). Asymmetry in the owl ear (both soft tissues and temporal parts of the skull, including modifications in the neurocranium and cartilaginous elements; Krings et al., 2020), serves to make the vertical directional sensitivity patterns different between the two ears for high frequencies, thus making possible vertical localization based on binaural comparison of intensity and spectral composition of sound (Norberg, 1977). Flatfishes (Pleuronectiformes) represent a diverse clade of benthic teleost fishes that possess a striking degree of cranial asymmetry exceeding that of any other vertebrate lineage (Black and Berendzen, 2020; Evans et al., 2021). The eyes of most flatfishes sit on the same side of the head; a development that happens during the larval stage where one eye of a symmetrical lava migrates to the other side. This produces a highly asymmetric cranium and has allowed the fish to colonise and dominate benthic aquatic habitats (Evans et al., 2021). Directional asymmetry is also widespread in invertebrates (see Okumura et al., 2008; Pélabon and Hansen, 2008), including wing size and shape (Klingenberg et al., 1998; Pélabon and Hansen, 2008), body shape (e.g., the Chilean magnificent beetle (*Ceroglossus chilensis*); Bravi and Benítez, 2013, and in the dextral spiralling of the shell in snails in the family Lymnaeidae; Okumuru et al., 2008), and eye morphology (e.g., cockeyed squids *Histioteuthis heteropsis* and *Stigmatoteuthis dofleini*; Thomas et al., 2017). Although it constitutes most of the examples in this study, directional asymmetry is by no means restricted to Animalia. Plants exhibit forms of asymmetry analogous to “handedness” in animals (Bahadur et al., 2019).

Simply, directional asymmetry is the natural condition for many taxa and avoiding the issue of left-right symmetry is not always an option if one wishes to accurately capture a specimen’s morphology. Here, we use toothed whales (odontocetes) as an example to firstly calculate how much directional asymmetry is present, and second, ensure that asymmetry is accurately quantified. Toothed whales (odontocetes) are well-known to have asymmetrical crania (Ness, 1967; Thompson, 1990; Fahlke et al., 2011; Churchill et al., 2018). This directional asymmetry is thought to have evolved as a result of an evolutionary hyperallometric investment into sound-producing soft tissue structures which have consequently driven evolution of the underlying bony structures, to facilitate high frequency vocalisation (echolocation) (Heyning and Mead, 1990). It is not entirely clear why the shift is sinistral (rather than dextral), but some propose it may be a by-product of selection pressure for a sinistrally positioned larynx and hyoid apparatus which provides a large, right piriform sinus whilst facilitating the swallowing of prey underwater (Macleod et al., 2007).

Examples such as these (and many others) underscore the apparent ubiquity of directional asymmetry in animals and plants, further bolstering our increasing knowledge of consistent left-right asymmetries (Klingenberg and Mclntyre, 1998a; Klingenberg et al., 2002). Although an increasingly well-known natural aspect of function and sometimes developmental morphology in some taxa, directional asymmetry is not often considered during standard geometric morphometric analyses and instead may be underrepresented (Fig.1). Further, previous studies have successfully addressed quantifying variation among individuals and asymmetry (Klingenberg et al., 1998; Klingenberg et al., 2002; Klingenberg, 2015), generally by calculating the Procrustes distance between a shape and its reflection, offering a measure of asymmetry (Bookstein 1991; Klingenberg and McIntyre 1998b; Mitteroecker and Gunz, 2009). The protocol proposed here builds on these previous methods with a focus on the benefits of using landmarks and semi-landmarks and, importantly, allowing the user to create a universal landmarking scheme regardless of a specimens’ asymmetry. This means symmetric and asymmetric specimens (for example from different species, genera, or families) can be analysed in the same geometric morphometric workflow.

Finally, the time-consuming nature of manually applying landmarks and semi-landmarks can impose limitations on data collection and project scope. For this reason, researchers use automated (Boyer et al., 2015a; 2015b) and semi-automated (Schlager et al. 2019) approaches to geometric morphometrics. However, these methods are not applicable to all morphologies. We estimate that using the method presented here reduces per-specimen processing time by about one third compared to applying semi-landmark curves to the whole specimen.

### Description

Standard geometric morphometric methods for mirroring landmarks and semi-landmarks (‘curves’) may misrepresent asymmetric specimens (Fig 1). Manually placing semi-landmarks on the entirety of an asymmetric specimen would provide a more accurate representation of the morphology; however, it is an extremely time-consuming solution. Here we offer a practical method for combining the two methods of manually landmarking and semi-landmarking asymmetric bones whilst mirroring bilaterally symmetrical bones. This method provides an accurate quantification of the morphology but also minimises the need and time taken to manually semi-landmark the entire specimen. We offer a solution for:

1. Quantifying asymmetry to determine whether landmarks should be manually placed rather than computer mirrored.
2. Using these data to create a landmarking protocol which will capture the morphology of both asymmetrical and symmetrical parts of the morphology, while minimising the time needed to semi-landmark the entire specimen (i.e., minimising per specimen processing time).
3. Producing an accurate representation of the morphology to then carry out standard geometric morphometrics.
4. Creating a pipeline that ensures both bilaterally symmetrical specimens and asymmetric specimens can be compared in the same analyses. The number and location of landmarks and semi-landmark curves is identical, *only the method of placement of landmarks is different*. This results in a global landmark and curve configuration that is the same among specimens regardless of whether they are asymmetrical or not, important for taxa for which some but not all specimens may have asymmetric morphologies.

Alternatively, we recommend that researchers looking at known bilaterally symmetrical specimens carry out step 1 (using landmarks only) to ascertain whether there is in fact some asymmetry or deformation in the specimen and thus whether quantifying asymmetry or reassessment of the specimen is required. If no asymmetry is detected, the researcher is assured that their specimen is bilaterally symmetric or does not have pronounced deformation and that mirroring of landmarks and semi-landmarks is sufficient to capture morphology.

## Materials and Methods

### Specimens and scan data collection

We demonstrate this approach using an example data set of odontocete (toothed whale) skulls. The data set comprises 157 odontocete skulls (Supporting Information: Table S1), representing 21 families, which range in asymmetry from no marked natural asymmetry to high levels of naso-facial asymmetry (see Coombs et al., 2020). Bilaterally symmetric mysticete (baleen whale) skulls are also used to visually illustrate the placement of curves on a bilaterally symmetrical specimen within the same analyses.

To analyse 3D geometric morphometric data that covers the entire cranium (to illustrate the point of full skull coverage of landmarks in asymmetric specimens) sampling was limited by specimen completeness and preservation. Inclusion of fossil specimens was determined by the extent of deformation and missing data. This does not mean that this method cannot be used on specimens with missing structures – see Supporting Information: Section 1: *Missing and variably present bones* to see how we dealt with incomplete specimens. Around 43% of specimens, including some extant specimens, had missing data, which was concentrated in the pterygoid, palate, jugal, squamosal, and tip of the rostrum (See Supporting Information: Section 1: *Missing and variably present bones*). Specimens with obvious taphonomic or other deformation were excluded from further analysis. Sexual dimorphism was not considered in this study as many fossils lack data on sex. All specimens are adults except for *Mesoplodon traversii* (NMNZ TMP012996) which is a sub-adult.

We scanned skulls using a Creaform Go!SCAN 20 or Creaform Go!SCAN 50 handheld surface scanner, depending on the size of the skull. Scans were initially cleaned, merged, and exported in ply format using VXElements v.6.0, and further cleaned and decimated in Geomagic Wrap software (3D Systems). We decimated models down to 1,500,000 triangles, reducing computational demands, while retaining sufficient detail for morphometric analysis. In many morphometric studies, it is possible to digitally reconstruct bilateral elements by mirroring across the midline plane if the skull (or object) is preserved on one side (Gunz et al., 2009; Gunz and Mitteroecker, 2013; Cardini et al., 2016 a, b). Due to a natural asymmetry occurring in the odontocete skull (Fahlke et al., 2011; Coombs et al., 2020), we limited mirroring to marginally damaged bones or easily mirrored missing bones only, where it was clear that mirroring would not mask asymmetric morphology. Elements were mirrored using the ‘mirror’ function in Geomagic Wrap (3D Systems). Skulls are used as the example throughout this study, but these methods could be used on any morphology as long as a midline is determined (see *Step 2: Quantifying asymmetry in the skull for details*).

This study focuses on how to capture the morphology of an asymmetric specimen. There are several general steps (i.e., not related to quantifying asymmetry specifically) that should be taken after *Step 4: Informing the curve protocol – which curves to manually place*, before mirroring curves. These additional steps have been highlighted here as a side note and are available in the detail in the Supporting Information: Section 1*: Additional steps before running geometric morphometric analyses* so as not to disrupt the ordering of the focus method of this paper; quantifying asymmetry.

These steps include:

- **Resampling**: As the placement of curves onto specimens is done manually points are likely not evenly placed along the bone. Curves are resampled to create even spacing between landmark points before being slid (see Supporting Information in Botton-Divet et al. 2016; Felice, 2020).
- **Sliding** the curves is the next crucial step, as equally spaced semi-landmarks should not (and cannot) be treated as optimally placed and the initial arbitrary placement of semi-landmarks can impose strong statistical artefacts (Gunz and Mitteroecker 2013; Bardua et al., 2019a).

These steps are not required for determining the placement of curves or in quantifying asymmetry but should be carried out once the curve protocol has been implemented. Code for these steps is provided in Felice (2020) and also via: https://github.com/EllenJCoombs/Quantifying_asymmetry

### Morphometric data collection – quantifying asymmetry

#### Step 1: Landmarking protocol

The placement of landmarks is the first step. These landmarks are then used to quantify asymmetry in the skull (or chosen morphology) (see *Step 2: Quantifying asymmetry*).

The first step is to quantify if, where, and how much asymmetry is evident in the skull. To do this, we placed 123 landmarks (sliding semi-landmarks are employed in a later step – see *Step 4: Informing the curve protocol*) irrespective of evidence of asymmetry over the entire surface of the skull (i.e., both sides) using Stratovan Checkpoint (Stratovan, Davis, CA, USA) (Fig. 2). We used the ‘single point’ option to add fixed landmarks. Landmarks were defined by Type I (biology) and Type II (geometry) (Bookstein, 1991; Bookstein, 1997) and were chosen to capture clearly homologous positions, e.g., tripartite sutures. Dentition was not landmarked. The landmark configuration for this data set is detailed in Fig.2 and Supporting Information; Table S2.

**Fig. 2.**
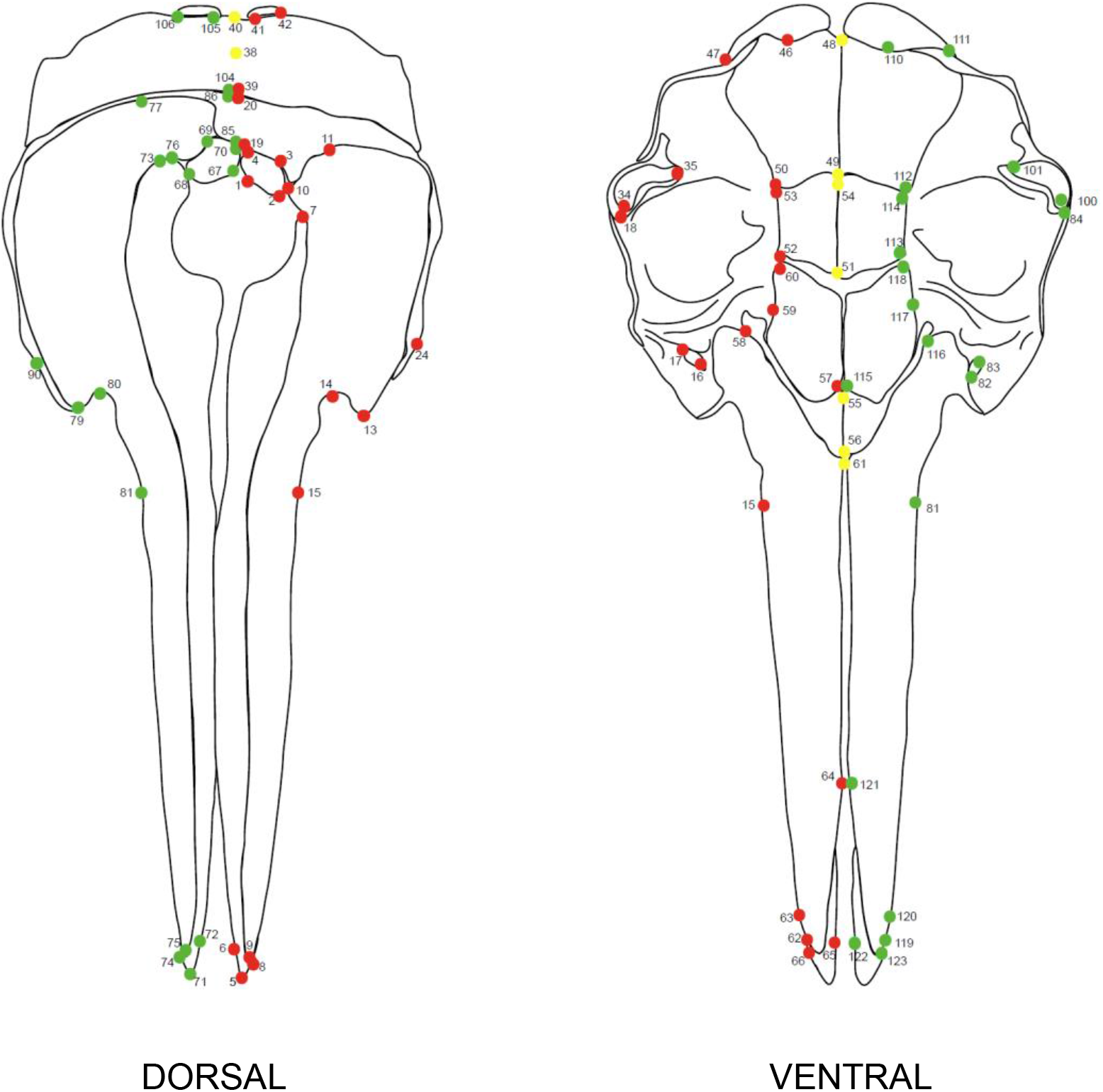
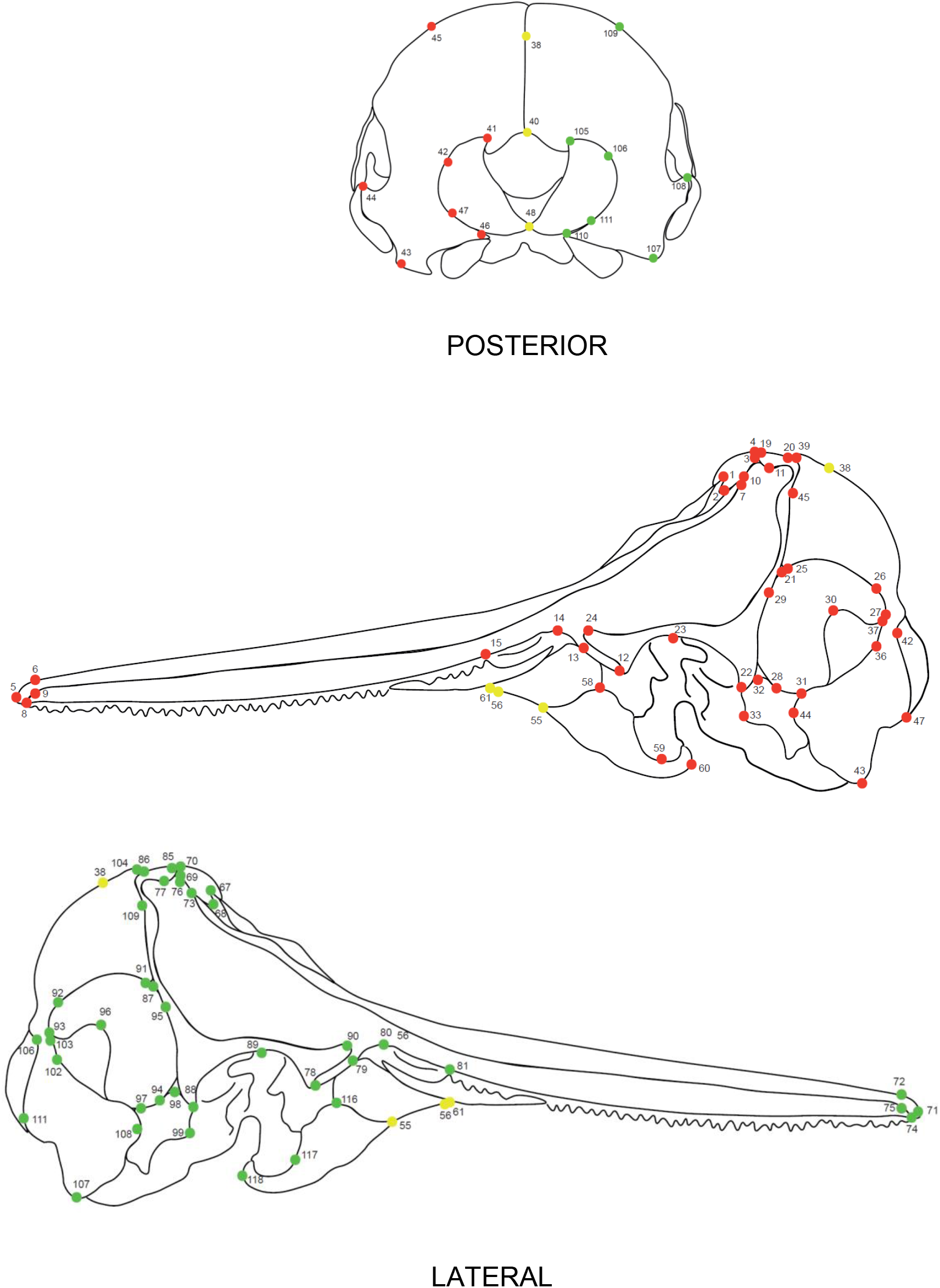
Landmark configuration on the cetacean skull. Red landmarks are placed on the left-hand side (LHS) of the specimen, with corresponding green landmarks also *manually* placed on the right-hand side (RHS) of the specimen. Midline landmarks are shown in yellow. Numbers correspond to descriptions shown in the Supporting Information (Table S2).

#### Step 2: Quantifying asymmetry in the skull

Once a whole skull landmarking protocol has been formulated and added to specimens, the next step is to quantify where asymmetry occurs in the skull or morphology. This is so that landmarks and curves can be manually placed on these regions, rather than being mirrored by an automated procedure (hereafter referred to as computer mirroring, as can be done with bilaterally symmetrical structures). To do this, the protocol to quantify asymmetry as in Coombs et al., 2020 is followed. A brief summary of the methods is provided here and the code for quantifying asymmetry is available in detail at: https://github.com/EllenJCoombs/Quantifying_asymmetry

1. Manually place landmarks on the entirety of the specimen (i.e., left and right sides, Fig. 2), as described in *Step 1*.
2. Generate mirrored landmarks for one side of the skull. To mirror the landmarks, we used 9 midline landmarks as an anchor (yellow landmarks, Fig. 2). We used the `mirrorfill` function in the R package paleomorph v.0.1.4 (Lucas and Goswami, 2017). NB. We decided to mirror the left-hand landmarks because of chirality in cetaceans, i.e., nasals are sinistrally shifted. This will be organism specific, and consideration must be taken to ensure specific morphologies are captured. In this example, mirroring the right-hand landmarks in cetaceans would exclude the nasals in some species. Consistency must be maintained between the side landmarks are mirrored from and to.
3. Compare the positions of the computer mirrored landmarks to those of the original, manually placed landmarks measuring the amount of landmark displacement between the two configurations.
4. Superimpose the specimens to remove all non-shape elements, i.e., size (scaling), translation, and rotation (positioning) from the data using Generalized Procrustes Analysis, here implemented in the gpagen function from the geomorph R package v.3.1.0 (Adams et al., 2019)
5. Calculate the Euclidean distances between a reference specimen (the computer-mirrored, landmarked specimen) (*Rn*) and a focal specimen (the manually landmarked specimen) (*Fn*). Both *Rn* and *Fn* are defined by three coordinates (*x*, *y*, *z*). The landmark displacements are measured for each landmark individually using the spherical coordinates system which measures between the *n*^th^ landmark of the *Fn* and the *Rn* specimens respectively, here implemented in the R package landvR v0.4 (Guillerme and Weisbecker, 2019).
6. If the specimen is asymmetric, the computer-mirrored landmark does not accurately reflect its morphology (Fig. 1). Estimate differences between *Fn* and *Rn* in the spherical coordinates system using the `coordinates.difference` function in landvR and extract the *ρ* (radius) for each landmark, for each specimen. This provides a measure of the Euclidean distance between a manually placed landmark which accurately represents the specimen’s morphology (*Fn)* and a computer -mirrored landmark (*Rn)*.
7. The larger the radii for a corresponding landmark the more displacement between *Fn* and *Rn.* We then interpret a higher *ρ* as an indication of more asymmetry in the skull (see Fig. 3 for a visualisation of this).

**Fig. 3.**
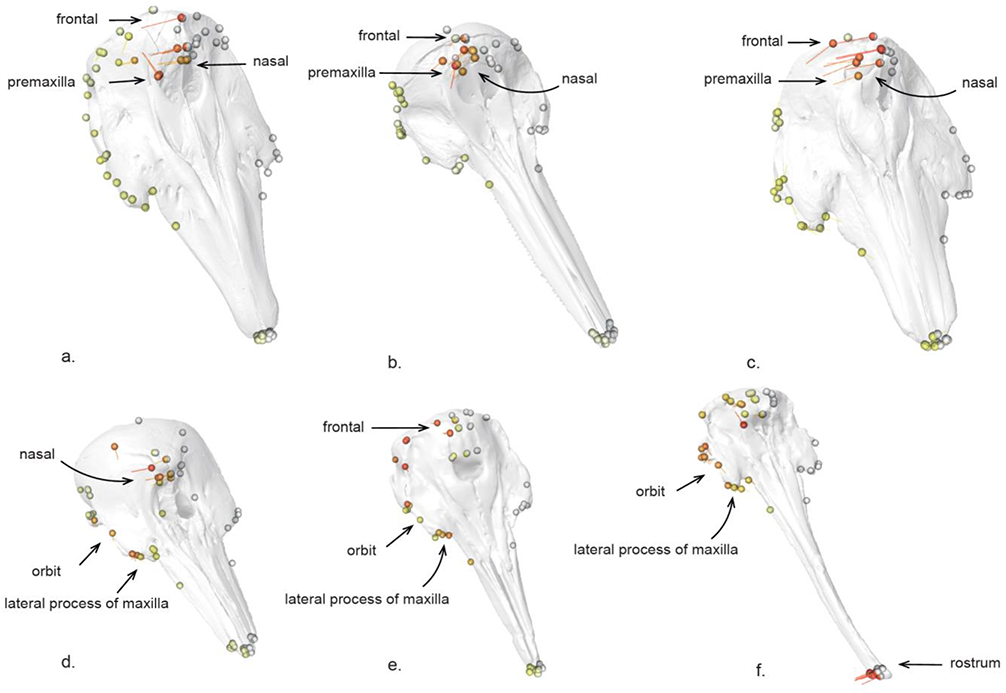
Visualisation of *p* (radii) from landvR showing asymmetry in the toothed whale skull. Landmarks are shown on a mesh of the skulls. The white spheres (landmarks) on the landvR outputs show the *fixed* landmarks (1–66) on the left-hand side (LHS) of the skull. The landmarks on the RHS of the skull vary in colour depending on how much difference there is between a computer-mirrored landmark (*Rn*) (which assumes the skull is bilaterally symmetrical) and a manually placed landmark (*Fn*) (which accurately depicts asymmetry). The larger the difference between the computer-mirrored landmark and the manually placed landmark, the hotter the colour. The highest amount of asymmetry is shown in red and dark orange, less asymmetry is shown in pale orange and yellow. The tails coming from each of the landmarks show how much and in which direction the landmarks have moved from where the computer mirrored them, to where the landmarks sit when manually placed. Specimens a-c show most asymmetry in the frontal, nasal, and dorsal, posterior premaxilla as is common in many odontocetes and is associated with echolocation (see Coombs et al., 2020 for details). Specimens d-f show areas of asymmetry in the nasal and frontal (d and e) but also in the orbit, lateral process of the maxilla, and the tip of the rostrum (f). Some ventral landmarks are shown to assist with visual interpretation – landmarks shown are dependent on the specimen and orientation of that skull for illustration of the method only. Specimens: a. *Delphinapterus leucas* (USNM 305071), b. *Delphinus delphis* (AMNH 75332), c. *Monodon monoceros* (USNM 267959), d. *Phocoena spinipinnis* (NHM 1900.5.7.29), e. *Pliopontos littoralis* (SAS 193), f. *Tagicetus joneti* (IRSNB/RBINS M.1892). Not shown to scale.

#### Step 3: Locating asymmetry in the skull or structure

Using landvR outputs for each of the specimens we can obtain a visual representation of where asymmetry occurs in the skull (Fig. 3) or structure. We recommend visualisation as a first step to ascertaining areas of asymmetry in the morphology. LandvR uses a ‘heat map’ approach to reflect displacement magnitude as shown in Fig. 3 (see Weisbecker et al., 2019 and Viacava et al., 2020 for further examples). Generally, we advise focusing on landmarks with the hottest colours (red, dark orange) at the least and investigating them further to a) check they are logical and b) obtain a numerical measure of the magnitude of asymmetry.

We can obtain a numerical value for asymmetry (i.e., displacement) by pulling out the radius value for each landmark and further calculating an average radius value for each landmark across the data set. This allows us to determine which landmarks exhibit the highest asymmetry. We then identify landmarks with high asymmetry for manual landmarking as they are the ones most likely to be misrepresented by mirroring alone. In this data set, the highest landmarks of variation are shown in Table 1. Fig. 3 shows the parts of the skull that were then considered for manual landmarking using the output from landvR.

**Table 1.**
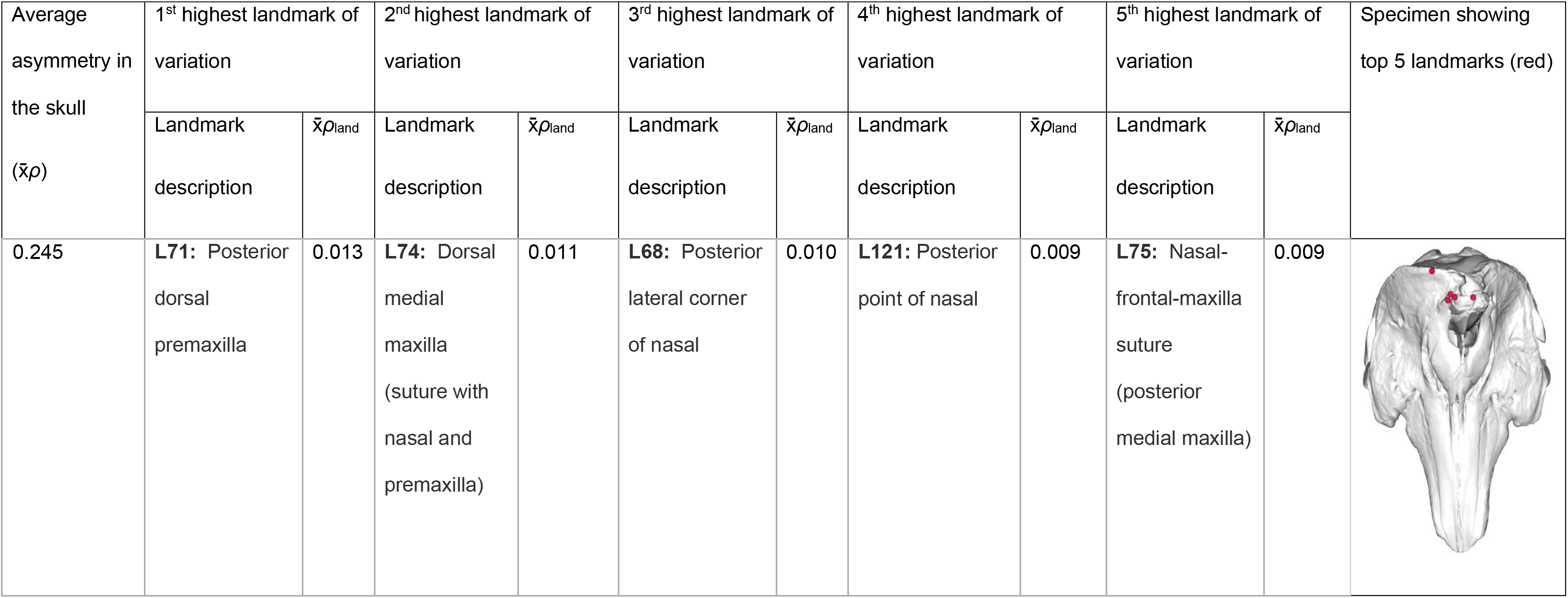
An example of the numerical outputs from landvR that help to inform areas of asymmetry in the skull or chosen morphology. Shown are the five landmarks with the greatest variation across the cranium for odontocetes in this study. x̄*ρ*_land_ is the average sum radii per landmark. Skull shown is *Monodon monoceros* (USNM 267959).

| Average asymmetry in the skull ($\bar{x}_\rho$) | 1<sup>st</sup> highest landmark of variation | | 2<sup>nd</sup> highest landmark of variation | | 3<sup>rd</sup> highest landmark of variation | | 4<sup>th</sup> highest landmark of variation | | 5<sup>th</sup> highest landmark of variation | | Specimen showing top 5 landmarks (red) |
| --- | --- | --- | --- | --- | --- | --- | --- | --- | --- | --- | --- |
| | Landmark description | $\bar{x}_{\rho_{land}}$ | Landmark description | $\bar{x}_{\rho_{land}}$ | Landmark description | $\bar{x}_{\rho_{land}}$ | Landmark description | $\bar{x}_{\rho_{land}}$ | Landmark description | $\bar{x}_{\rho_{land}}$ | |
| 0.245 | **L71:** Posterior dorsal premaxilla | 0.013 | **L74:** Dorsal medial maxilla (suture with nasal and premaxilla) | 0.011 | **L68:** Posterior lateral corner of nasal | 0.010 | **L121:** Posterior point of nasal | 0.009 | **L75:** Nasal-frontal-maxilla suture (posterior medial maxilla) | 0.009 |  |

To investigate the landmarks of highest variation (and thus potential candidates for manual placement) we extract the ‘radii’ which is the radius per landmark for each specimen and ‘`radii_mean`’ which is the mean radius per landmark. An example is provided below. Details are provided on Github (https://github.com/EllenJCoombs/Quantifying_asymmetry). An example of the results you can obtain using the below code are shown in Table 1. Much more can be done with these results should the user wish to investigate and visualise mean shapes and Procrustes distance, to name just two possibilities. See Guillerme and Weisbecker (2019) for further code and visualisations.

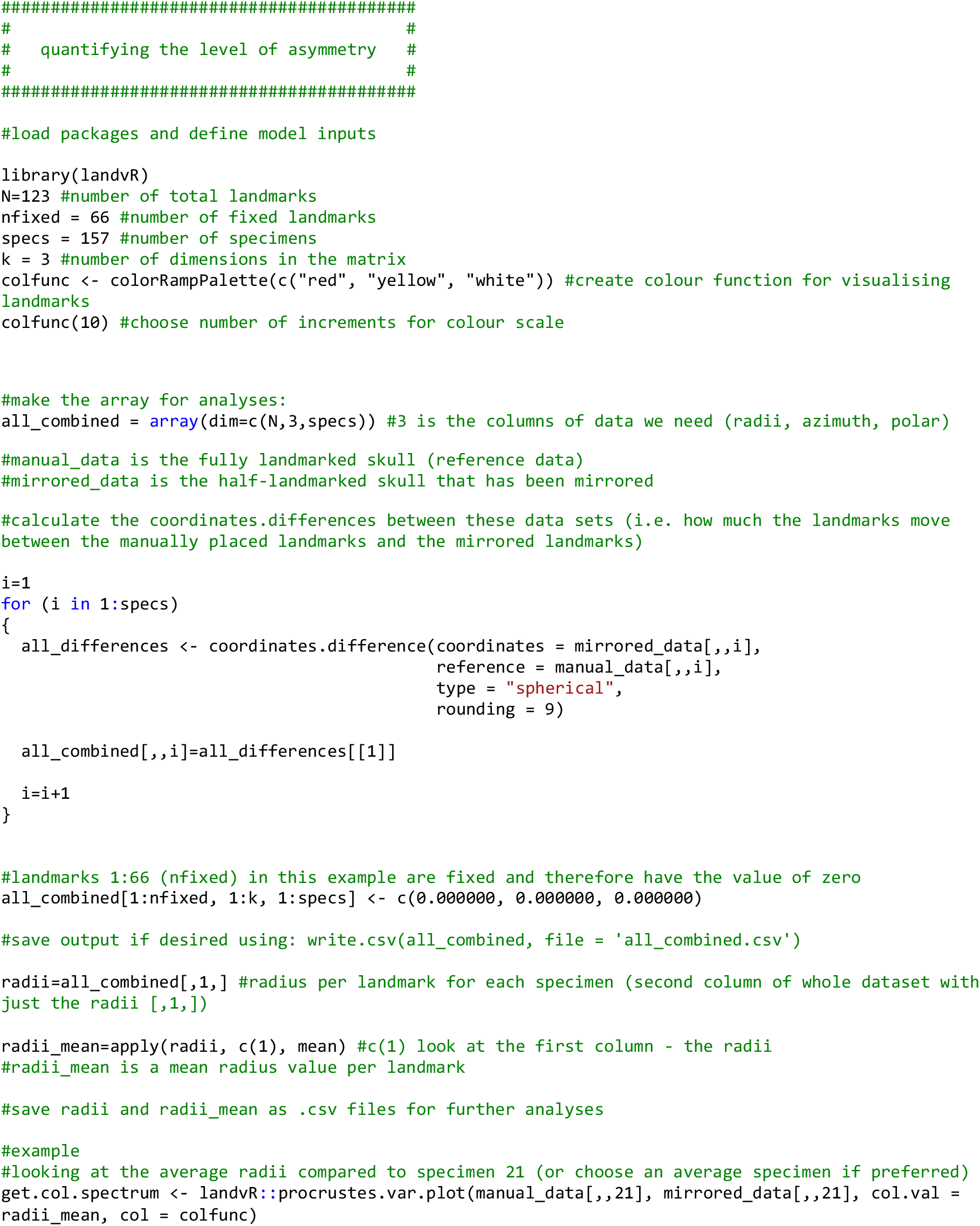

**Code snippet 1**: Extracting the radii and average radii for each specimen using the landvR package

#### Step 4: Informing the curve protocol – which curves to manually place

For this data set, the top landmarks of variation were concentrated in the nasals, frontal, premaxilla, and maxilla (Fig. 3, Table 1). Asymmetry was also found in the orbit, lateral process of the maxilla, and the tip of the rostrum (Fig. 3 d-f). This informs our protocol for manually placing landmarks and curves on these bones, instead of mirroring. It also informs the landmarks and curves which can be mirrored, i.e., those that showed little asymmetry (pale yellow) such as the ventral and posterior of the skull in this example. Curves are then manually placed on one side of the skull for symmetrical structures (as is standard in bilaterally symmetrical specimens), with the addition of manual placement of curves on both sides of the face (maxilla, premaxilla, nasals, and frontal) to capture the morphology of asymmetric bones (Fig. 4). See Supporting Information; Table S3 for curve information.

**Fig. 4.**
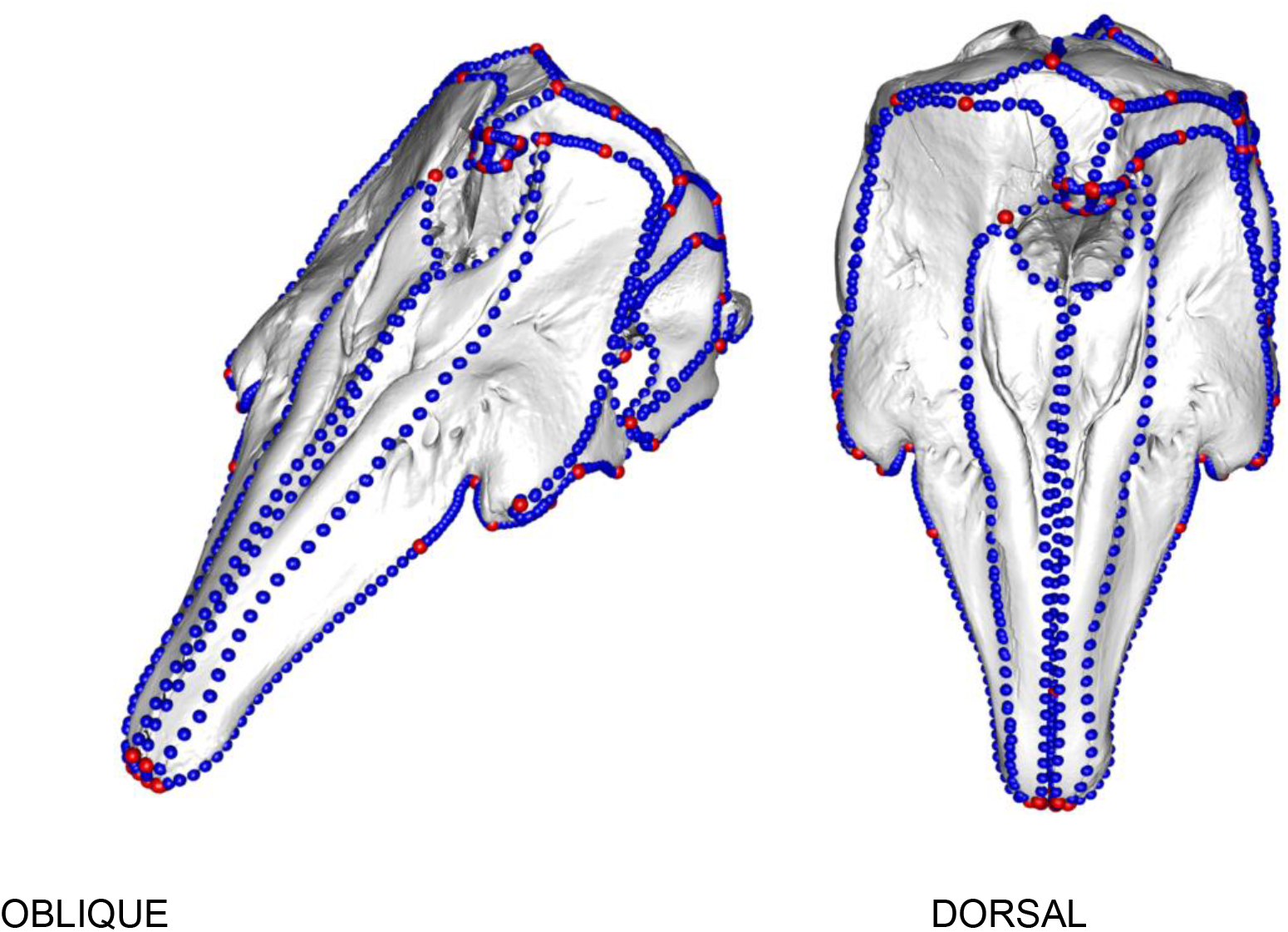
Curves and landmarks manually placed on the asymmetric cetacean face. Note the asymmetry in the posterior of the skull near the nasal. The manual placement of landmarks on the right hand side (RHS) of the skull are selected based on the results from landvR (Table 1, Fig. 3). Note the RHS posterior and the ventral skull are not shown with curves here because these are bilaterally symmetrical parts of the morphology (smaller radii values from landvR on the posterior and ventral of the skull) and thus landmarks and semi-landmark curves can be computer mirrored onto these sections (see Fig. 5 for full skull details). The skull on the left is shown in oblique view, the skull on the right is shown in dorsal view. Specimen shown is *Delphinapterus leucas* (USNM 305071).

See the methods section for infromation on resampling and sliding curves. This step is not specific to the asymmetric protocol that we address in this study but would be carried out at this stage, before mirroring curves.

#### Step 5: Placing landmarks and semi-landmarks on asymmetrical specimens

We use the results from landvR to determine which of the bones in the skull were asymmetric and thus requiring manual landmarking and which could be reliably placed by mirroring bilaterally symmetric landmarks across the skull midline. For the asymmetric specimens, we placed 57 landmarks on the LHS of the skull and nine landmarks on the midline. We mirrored 33 landmarks to symmetrical bones on the right-hand side (RHS) of the skull and we manually placed 24 landmarks on asymmetric bones on the RHS of the skull. We manually placed 60 curves using the ‘curve’ option in Checkpoint, on the sutures between bones on the LHS of the skull and four curves on the midline (Fig. 5). We manually placed 21 curves on asymmetrical bones (the face) on the RHS, mostly concentrated in the nasals, dorsal premaxilla, dorsal maxilla, orbit and rostrum, and the rest were computer-mirrored from the LHS (Fig. 5). Curves should then be resampled and slid (see Methods).

**Fig. 5.**
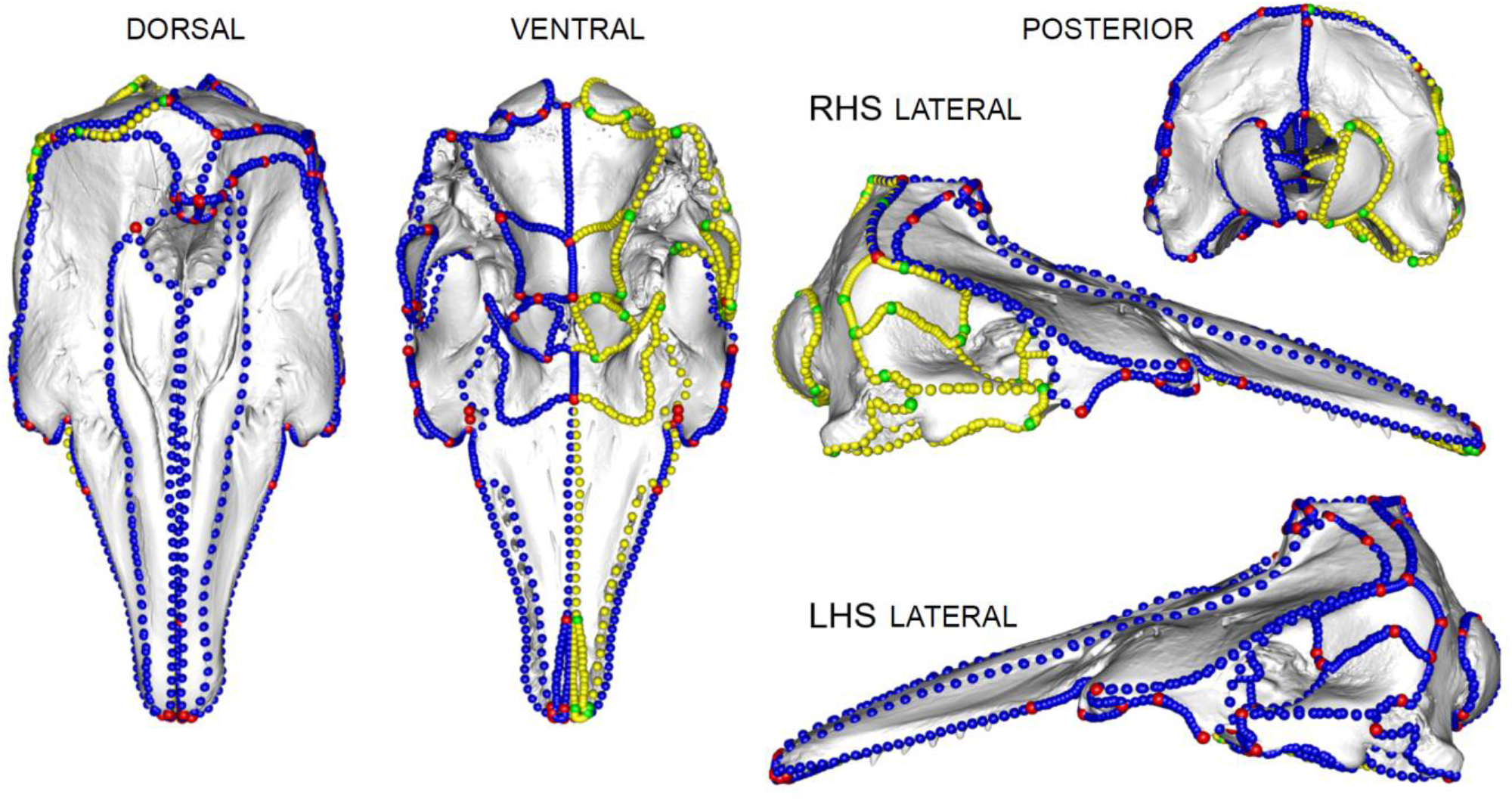
Landmark configuration on an asymmetric skull. Red = manually placed landmarks on asymmetric bones, green = computer mirrored landmarks on symmetric bones, blue = manually placed curves (semi-landmarks) on asymmetric bones, yellow = computer mirrored curves (semi-landmarks) on symmetric bones. Specimen is *Delphinapterus leucas* (USNM 305071).

Code snippet 2 shows how to mirror bilaterally symmetrical curves only, whilst leaving manually placed (asymmetric) curves untouched. This results in manually placed asymmetric curves and computer mirrored bilaterally symmetrical curves being combined to cover the entire skull or morphology (Fig. 5). Using this method (code snippet 2) ensures that both bilaterally symmetrical specimens and asymmetric specimens (for example if specimens in the sample, e.g., a specific species, sex, or developmental stage have asymmetry, but other specimens do not) can still be compared in the same analyses as the number and location of landmarks and semi-landmark curves are identical, *only the method of placement is different*.

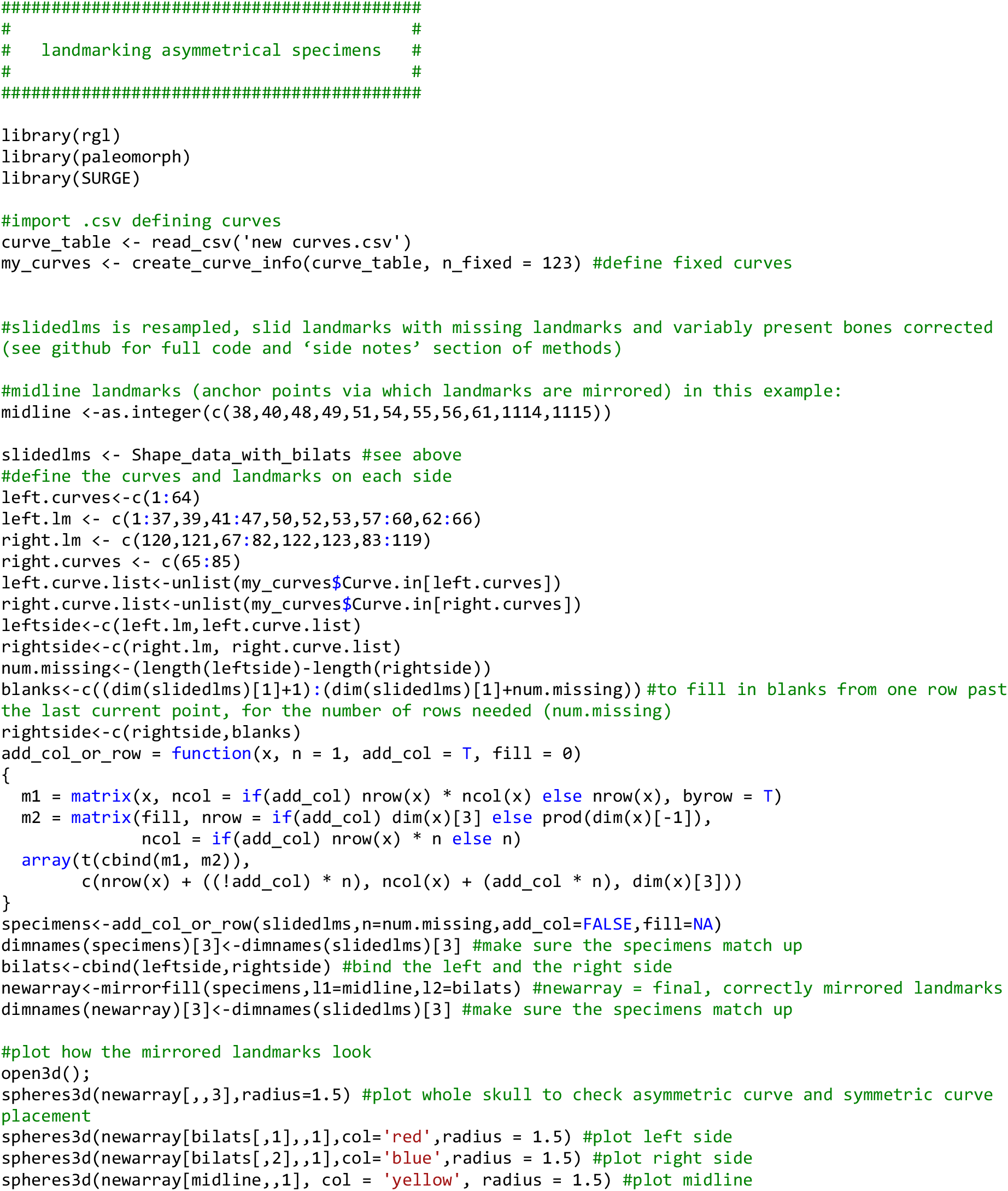

**Code snippet 2**: Code for mirroring bilaterally symmetrical curves alongside manually placed asymmetric curves. Complete code for mirroring symmetrical specimens is available at: https://github.com/EllenJCoombs/Quantifying_asymmetry

## Results

### Landmark and curve placement on asymmetrical specimens

Once these steps are followed, the user will have quantified where asymmetry exists in the specimen(s) and created a curve protocol that not only captures asymmetry in the structure or specimen but also accounts for any bilateral symmetry if present in that same structure or in other specimens in the data set. Examples of successful placement of curves in asymmetric and symmetric specimens are shown here (Fig. 5-7).

### Additionally: Mirroring landmarks on bilaterally symmetrical specimens

On symmetrical specimens, here represented by the mysticetes (baleen whales) (see Fahlke and Hampe, 2015; Coombs et al., 2020) (Fig. 6), we placed 57 landmarks on the left-hand side (LHS) of the skull and nine landmarks on the midline. We placed 60 sliding semi-landmark curves on the sutures between bones on the LHS of the skull and four curves on the midline. These curves and landmarks were then mirrored (using the midline landmarks and curves as an anchor) using the `mirrorfill` function in the R package ‘paleomorph’ v.0.1.4 (See: https://github.com/EllenJCoombs/Quantifying_asymmetry). This method (see code snippet 2) ensures that both bilaterally symmetrical specimens and asymmetric specimens can be compared in the same analyses as landmark and curve numbers match between specimens. This results in a global landmark and curve configuration that is the same among specimens regardless of whether they are asymmetrical or not (Fig. 7).

**Fig. 6.**
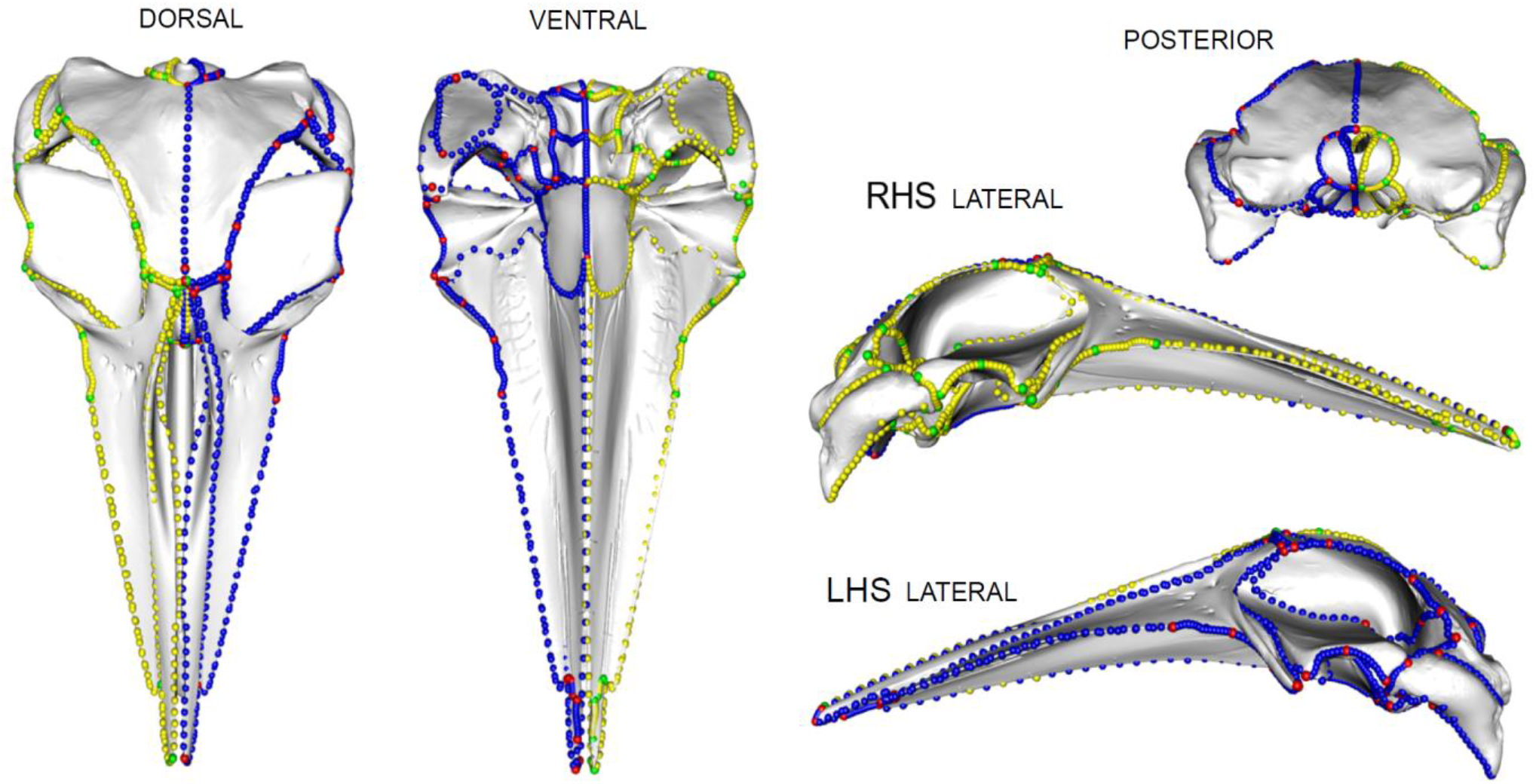
Landmark configuration on a symmetric skull. Landmark protocol for the symmetric mysticete. Red = manually placed landmarks, green = computer mirrored landmarks, blue = manually placed curves (semi-landmarks), yellow = computer mirrored curves (semi-landmarks). Specimen is *Balaenoptera acutorostrata* (NHM 1965.11.2.1).

**Fig 7.**
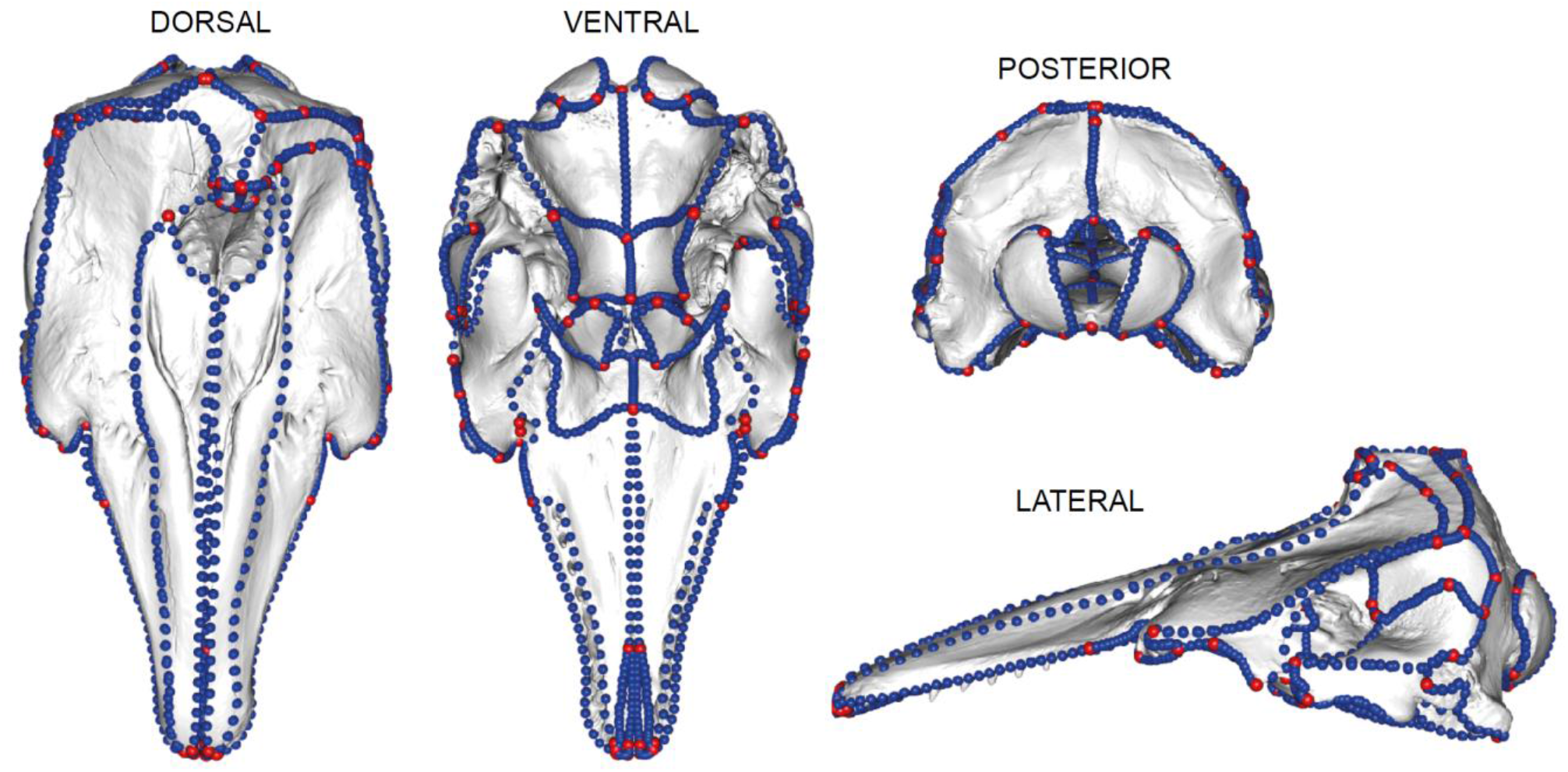
Final landmark and curve sliding semi-landmark placement on all skulls regardless of asymmetry. The landmarks in red are type I and type II landmarks. The curves in blue define outlines and margins of bones. There are 123 landmarks and 124 curves on this specimen. Landmarks and curves shown on a beluga (*Delphinapterus leucas* (USNM 305071)) specimen. The methods of placement of these landmarks and curves are different depending on whether the specimen is bilaterally symmetrical or asymmetrical; however, the finished result (i.e., number of landmarks and curves and placement on bones) is uniform across all specimens so that morphology is comparable.

Some odontocetes, such as phocoenids show little asymmetry in the skull (Cranford et al., 1996; Marx et al., 2016). However, others, particularly highly asymmetrical specimens such as the kogiids and physeteroids could be misrepresented in the morphospace when using computer landmarks. In the example below we see that asymmetric species (circled; Fig. 8) do shift in morphospace position if landmarks are mirrored or manually placed. This is particularly evident in asymmetric specimens such as monodontids, kogiids (Ness, 1967), and Physeteroidea (Coombs et al., 2020).

**Fig. 8.**
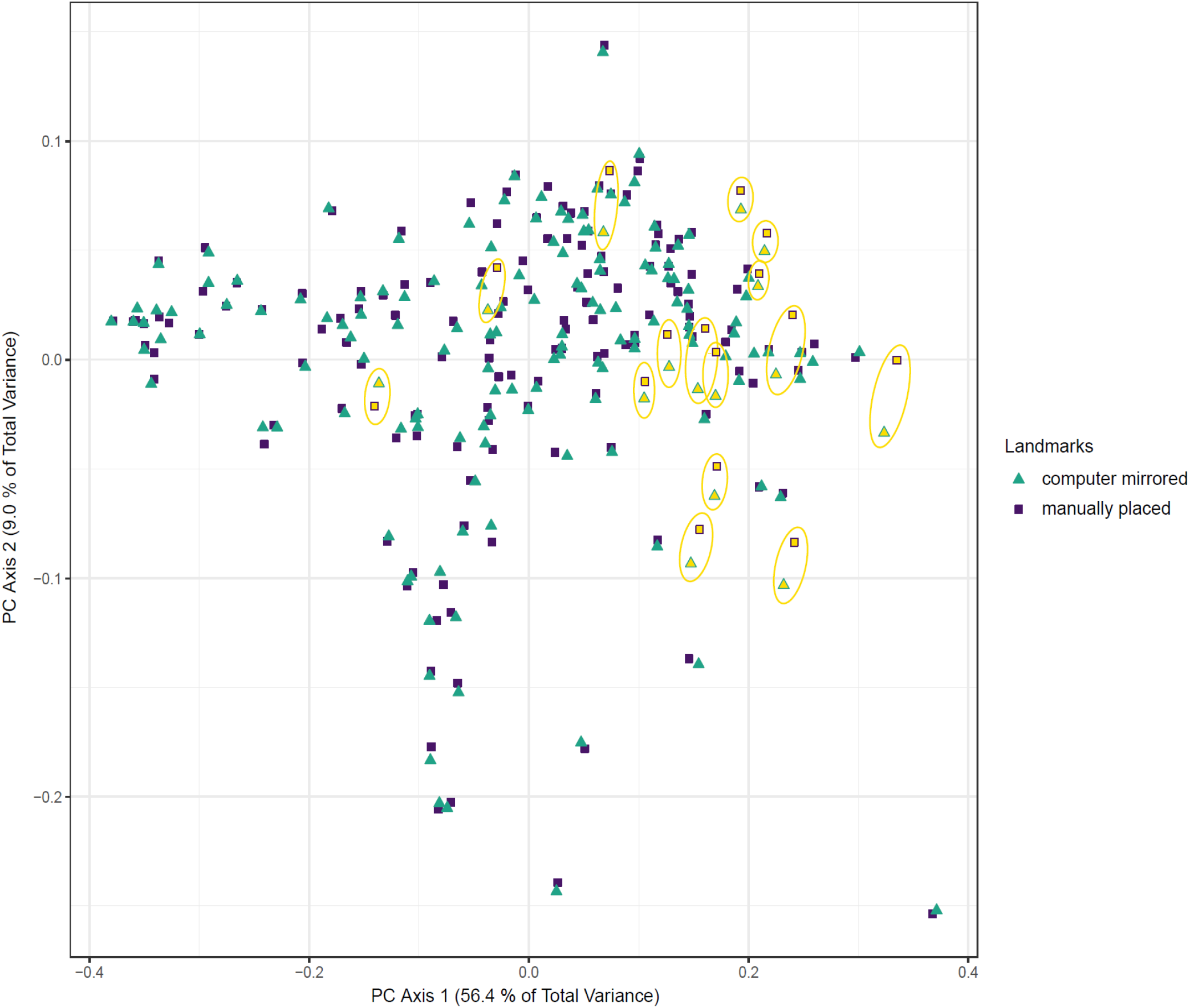
An example of how specimens with some asymmetry (here concentrated in the naso-facial region of the skull) will sit in the morphospace when landmarked manually or using computer mirrored landmarks. The specimens with the greatest difference between their computer mirrored landmarks and manually placed landmarks (circled in yellow) are those with higher asymmetry. Morphospace of 157 odontocete skulls with landmark and semi-landmarks over the entirety of the skull (both asymmetric and symmetric bones).

Finally, for this specific data set, landmarking a skull using the method presented here took the researcher around a third less time than manually landmarking the whole skull.

## Discussion

Directional asymmetry in organisms is a fascinating phenomenon but can complicate data collection by making automated mirroring of morphometric data inappropriate. Methods such as mirroring landmarks is a standard technique used for bilaterally symmetrical specimens (Gunz et al., 2009; Cardini et al., 2010; Gunz and Mitteroecker, 2013) which provide an accurate quantification of morphology whilst reducing the time needed to landmark the entire surface of the skull (Bardua et al., 2019a). However, while this is a suitable technique for bilaterally symmetrical structures, it may misrepresent asymmetric structures. Further, it can be difficult to detect asymmetry ‘by eye’ and then landmark a specimen as such based on a visual interpretation. Not accurately quantifying asymmetry in the initial stages could result in asymmetry being missed or perhaps even overrepresented. Here we offer a practical method to accurately quantify the morphology but also minimise the time by about a third (in this example data set) to manually landmark the entire specimen.

Natural, directional asymmetry occurs when one side of the structure is consistently different (e.g., in size or shape) (Graham et al., 1993; Parés-Casanova, 2020). Directional asymmetry is recorded in many specimens, from algae and leaf blades to corals, turtles, and birds. In some taxa, asymmetry can be genus or even sex specific, for example, male speckled wood butterflies (*Pararge aegeria*) have directional asymmetry in the fore and hindwing and forewing width (Windig and Nylin, 1999) and in cetaceans, families such as the monodontids and the kogiids have more naso-facial asymmetry than families such as the delphinids (Ness, 1967). It is therefore useful to have a protocol that can be used to capture morphology in both asymmetric and symmetric specimens that are to be analysed together. This protocol results in a global landmark and curve configuration that is the same among specimens regardless of whether they are asymmetrical or not (Fig. 6; Fig. 7).

This protocol provides a substantial increase in data collection speed. The time-consuming nature of digitising 3D landmark and semi-landmark data can impose limitations on sampling in resource-limited research projects. This has led some researchers to seek automated (Boyer et al., 2015a; 2015b) or semi-automated (Schlager et al. 2019) approaches to geometric morphometrics, but these methods are not applicable to all morphologies or hypotheses. We estimate that using the method presented here (i.e., manually semi-landmarking asymmetric bones and mirroring semi-landmarks for bilaterally symmetric bones only) reduces per-specimen processing time by about one third compared to applying semi-landmark curves to the whole skull. Gains in digitisation speed will be specific to the data set in question, for example, taxa with more asymmetry would require more manual landmarking and thus an increased time investment to accurately capture the asymmetric morphology.

A time effective method is desirable to any researcher but most important is the accurate representation of a specimen and its morphology. This method helps to first quantify any asymmetry in the morphology, and then to accurately represent it. In the example shown (Fig. 8) this goes as follows; computer mirrored landmarks and curves and manually placed landmarks and curves are placed on odontocete skulls to observe the difference that incorrectly placed landmarks can have on reporting morphology. Importantly, the difference between manual and mirrored specimens can be as great as the difference between species (Fig. 8) and thus has the potential to mislead downstream analyses. We find that the incorrect placing of specimens in the morphospace (via incorrect landmarking) can place specimens as far from their true morphology (if correctly landmarked) as from other species. This in turn could influence results that may be looking at, for example, ecological influences on morphology, such as species-specific diet, habitat, or other ecological factors. It is therefore central to ascertain which specimens in a sample could be misrepresented by mirroring of landmarks.

### Limitations

Finally, there are some limitations to this exploratory approach. Firstly, the method is most likely useful for studies where measurement error is small compared to biological variation, for example, macroevolutionary studies, interspecific studies, ontogenetic studies, and studies of sexual dimorphism. Variation in intraspecific studies may be small and difficult to quantify using this method. That said, we still recommend quantifying asymmetry in intraspecific cohorts and on specimens with assumed bilateral symmetry if only to a) confirm the latter and thus support the mirroring of landmarks from one half of the morphology to the other, or b) highlight any deformation in specimens, especially fossils. Secondly, an understanding of the specimen’s morphology is desirable to interpreting the outputs from landvR, for example, it is useful to know whether landmarks that show up in hotter colours (red, dark orange) are reflective of the biology or an artefact of deformation. An in-depth anatomical knowledge of study specimens is not a prerequisite, but we do recommend considering asymmetric landmarks carefully to ascertain whether any observed asymmetry is likely biological or deformational.

The code for these analyses is available at: https://github.com/EllenJCoombs/Quantifying_asymmetry. The code relies heavily on functions available in the SURGE package (Felice, 2020) and the Paleomorph package (Lucas and Goswami, 2017). Due to advances in coding and imaging technologies, we anticipate continual updates to these methods and welcome user suggestions and contributions.

## Acknowledgements

We would like to thank Anjali Goswami for her comments on drafts of this manuscript and Thomas Guillerme for help and assistance with the landvR package. Thanks also go to the many curators and museum staff who helped EJC collect scan data.

## Author contributions

EJC conceived the idea. EJC and RNF designed the methodology. EJC collected and analysed the data. RNF compiled code. Both authors contributed critically to the drafts and gave final approval for publication.

## Data availability

All data and code are freely available and are stored at: https://github.com/EllenJCoombs/Quantifying_asymmetry

## Supporting information

Additional supporting information may be found in the Supporting Information section.

