## Supporting information for "Quantifying asymmetry in non-symmetrical morphologies, with an example from Cetacea"

Ellen. J. Coombs^1,2^

Ryan. N. Felice2^2,3^

1. Department of Genetics, Evolution, and Environment, University College London, London, UK
2. Department of Life Sciences, The Natural History Museum, London, UK
3. Centre for Integrative Anatomy, Department of Cell and Developmental Biology, University College London, London, UK

### **Table S1.** Specimen list in alphabetical order. Additional details on family and suborder.

| Museum ID | Family | Suborder |
| --- | --- | --- |
| *Acrodelphis* UCMP 73695 | Kentriodontidae | odontocete |
| *Agorophiid* USNM 205491 | Agorophiidae | odontocete |
| *Albertocetus* ChM PV8680 | Xenorophidae | odontocete |
| *Albireo whistleri* UCR 14589 | Albireonidae | odontocete |
| *Ankylorhiza tiedemani* CCNHM 103 | incertae sedis | odontocete |
| *Aprixokogia kelloggi* USNM 187015 | Kogiidae | odontocete |
| *Argyrocetus joaquinensis* USNM 11996 | Delphinida | odontocete |
| *Atocetus iquensis* MNHN.F.PPI. 113 | Kentriodontidae | odontocete |
| *Aulophyseter morricei* UCMP 81661 | Physeteridae | odontocete |
| *Berardius arnuxii* NHM 1935.10.23.1 | Ziphiidae | odontocete |
| *Berardius bairdii* NHM 1954.9.21.1 | Ziphiidae | odontocete |
| *Berardius mimimus* USNM 276375 | Ziphiidae | odontocete |
| *Brachydelphis mazeazi* MUSM 564 | Pontoporiidae | odontocete |
| *Cephalorhynchus commersonii* NN | Delphinidae | odontocete |
| *Cephalorhynchus eutropia* NHM 1881.8.17.1 | Delphinidae | odontocete |
| *Cephalorhynchus heavisidii* NHM 1948.7.27.1 | Delphinidae | odontocete |
| *Cephalorhynchus hectori maui* NMNZ MM002607 | Delphinidae | odontocete |
| *Cephalorhynchus hectori* NMNZ MM002288 | Delphinidae | odontocete |
| *Chavinziphius maxillocristatus* MUSM 2538 | Ziphiidae | odontocete |
| *Chilcacetus cavirhinus* MUSM 1401 | incertae sedis | odontocete |
| *Cotylocara macei* CCNHM 101 | Xenorophidae | odontocete |
| *Delphinapterus leucas* USNM 305071 | Monodontidae | odontocete |
| *Delphinodon dividum* USNM 7278 | Kentriodontidae | odontocete |
| *Delphinus capensis* NHM 1981.7.11 | Delphinidae | odontocete |
| *Delphinus delphis* AMNH 75332 | Delphinidae | odontocete |
| *Dilophodelphis fordycei* USNM 214911 | Platanistidae | odontocete |
| *Echovenator sandersi* GSM 1098 | Xenorophidae | odontocete |
| *Ensidelphis riveroi* MUSM 3898 | incertae sedis | odontocete |
| *Etruridelphis* PU 13884 | Delphinidae | odontocete |
| Eurhinodelphid UCMP 99669 | Eurhinodelphinidae | odontocete |
| *Eurhinodelphinidae chilcocetus* MUSM | Eurhinodelphinidae | odontocete |
| *Eurhinodelphis cocheteuxi* IRSNB | Eurhinodelphinidae | odontocete |
| *Eurhinodelphis longirostris* USNM 244404 | Eurhinodelphinidae | odontocete |
| *Feresa attenuata* USNM 504916 | Delphinidae | odontocete |
| *Globicephala macrorhynchus* NHM 1912.10.27 | Delphinidae | odontocete |
| *Globicephala melas* NMNZ MM001946 | Delphinidae | odontocete |
| *Globicephala sp* USNM 21867 | Delphinidae | odontocete |
| *Goedertius oregonensis* LACM 123887 | Allodelphinidae | odontocete |
| *Grampus griseus* USNM 571602 | Delphinidae | odontocete |
| *Hemisyntrachelus cortesii* MBGPT NN | Delphinidae | odontocete |
| *Hemisyntrachelus oligodon* SMNK-PAL 3841 | Delphinidae | odontocete |
| *Huaridelphis raimondii* MUSM 1396 | Squalodelphinidae | odontocete |
| *Hyperoodon ampullatus* NHM 1992.42 | Ziphiidae | odontocete |
| *Hyperoodon planifrons* NHM 1952.9.30.1 | Ziphiidae | odontocete |
| *Indopacetus pacificus* USNM 593534 | Ziphiidae | odontocete |
| *Inia geoffrensis* AMNH 93415 | Iniidae | odontocete |
| *Kampholophus serrulus* UMCP 36045 | Kentriodontidae | odontocete |
| *Kentriodon pernix* USNM 10670 | Kentriodontidae | odontocete |
| *Kentriodon schneideri* USNM 323772 | Kentriodontidae | odontocete |
| *Kogia breviceps* USNM 22015 | Kogiidae | odontocete |
| *Kogia simus* NHM.1952.8.28.1 | Kogiidae | odontocete |
| *Koristocetus pescei* MUSM 888 | Kogiidae | odontocete |
| *Lagenodelphis hosei* USNM 571619 | Delphinidae | odontocete |
| *Lagenorhynchus acutus* USNM 504196 | Delphinidae | odontocete |
| *Lagenorhynchus albirostris* AMNH 37162 | Delphinidae | odontocete |
| *Lagenorhynchus australis* 1944.11.30.1 | Delphinidae | odontocete |
| *Lagenorhynchus cruciger* NHM 1960.8.24.1 | Delphinidae | odontocete |
| *Lagenorhynchus obliquidens* NHM 1992.83 | Delphinidae | odontocete |
| *Lagenorhynchus obscurus* NHM 1846.3.11.8 | Delphinidae | odontocete |
| *Lamprolithax simulans* LACM 37858 | incertae sedis | odontocete |
| *Lipotes vexillifer* AMNH 57333 | Lipotidae | odontocete |
| *Lissodelphis borealis* USNM 550188 | Delphinidae | odontocete |
| *Lissodelphis peronii* NMNZ MM002116 | Delphinidae | odontocete |
| *Livyatan melvillei* MSNUP | incertae sedis | odontocete |
| *Lomacetus ginsburgi* MNHN.F.PPI.104 | Phocoenidae | odontocete |
| *Macrokentriodon* CMM V 15 | Kentriodontidae | odontocete |
| *Macrosqualodelphis ukupachai* MUSM 2545 | Squalodelphinidae | odontocete |
| *Mesoplodon bidens* USNM 593438 | Ziphiidae | odontocete |
| *Mesoplodon bowdoini* NMNZ MM001900 | Ziphiidae | odontocete |
| *Mesoplodon carlhubbsi* USNM 504128 | Ziphiidae | odontocete |
| *Mesoplodon densirostris* NMV C 36362 | Ziphiidae | odontocete |
| *Mesoplodon europaeus* USNM 571665 | Ziphiidae | odontocete |
| *Mesoplodon ginkgodens* USNM 298237 | Ziphiidae | odontocete |
| *Mesoplodon grayi* USNM 49880 | Ziphiidae | odontocete |
| *Mesoplodon hectori* NHM 1949.8.19.1 | Ziphiidae | odontocete |
| *Mesoplodon hotaula* USNM 593426 | Ziphiidae | odontocete |
| *Mesoplodon layardii* USNM 550150 | Ziphiidae | odontocete |
| *Mesoplodon mirus* USNM 504612 | Ziphiidae | odontocete |
| *Mesoplodon perrini* LACM 97501 | Ziphiidae | odontocete |
| *Mesoplodon peruvianus* USNM 571258 | Ziphiidae | odontocete |
| *Mesoplodon stejnegeri* USNM 504330 | Ziphiidae | odontocete |
| *Mesoplodon traversii* juv. NMNZ TMP012996 | Ziphiidae | odontocete |
| *Messapicetus gregarius* MUSM 1481 | Ziphiidae | odontocete |
| *Messapicetus longirostris* MSNUP | Ziphiidae | odontocete |
| *Monodon monoceros* USNM 267959 | Monodontidae | odontocete |
| MUSM 563 | Lophocetinae | odontocete |
| MUSM 605 | Lophocetinae | odontocete |
| *Nazcacetus urbinai* MUSM 949 | Ziphiidae | odontocete |
| *Neophocaena asiaeorientalis* USNM 240001 | Phocoenidae | odontocete |
| *Neophocoena phocaenoides* NHM 1903.9.12.3 | Phocoenidae | odontocete |
| *Notocetus vanbenedeni* MUSM 1395 | Squalodelphinidae | odontocete |
| *Odobenocetops peruvianus* SMNK PAL 2491 | Odobenocetopsidae | odontocete |
| *Orcaella brevirostris* NHM.1883.11.20.2 | Delphinidae | odontocete |
| *Orcaella heinsohni* USNM 284430 | Delphinidae | odontocete |
| *Orcinus orca* USNM 11980 | Delphinidae | odontocete |
| *Orycterocetus crocodilinus* USNM 22926 | Physeteridae | odontocete |
| *Papahu taitapu* OU 22066 | Waipatiidae | odontocete |
| *Parapontoporia sternbergi* SDNHM 75060 | Lipotidae | odontocete |
| Patriocetid new genus ChM PV4753 | Patriocetidae | odontocete |
| Patriocetid or Waipatiid new genus CCNHM 1078 | Patriocetidae | odontocete |
| *Patriocetus ehrlichii* 1999-3 Cet. 4 | Patriocetidae | odontocete |
| *Patriocetus sp* MB Ma. 42882 | Patriocetidae | odontocete |
| *Peponocephala electra* USNM 504511 | Delphinidae | odontocete |
| *Phocoena dioptrica* NHM 1939.9.30.1 | Phocoenidae | odontocete |
| *Phocoena phocoena* AMNH 212161 | Phocoenidae | odontocete |
| *Phocoena sinus* SDNHM 20697 | Phocoenidae | odontocete |
| *Phocoena spinpinnis* NHM 1900.5.7.29 | Phocoenidae | odontocete |
| *Phocoenoides dalli* USNM 276062 | Phocoenidae | odontocete |
| *Physeter macrocephalus* NHM 2007.1 | Physeteridae | odontocete |
| *Piscolithax longirostris* SAS 933 | Phocoenidae | odontocete |
| *Piscolithax tedfordi* UCMP 15972 | Phocoenidae | odontocete |
| *Platanista gangetica* USNM 172409 | Platanistidae | odontocete |
| *Pliopontos littoralis* SAS 193 | Pontoporiidae | odontocete |
| *Pomatodelphis* CMM V 3915 | Platanistidae | odontocete |
| *Pomatodelphis* USNM 187414 | Platanistidae | odontocete |
| *Pontoporia blainvillei* USNM 482727 | Pontoporiidae | odontocete |
| *Prosqualodon davidis* USNM 467596 | Prosqualodontidae | odontocete |
| *Pseudorca crassidens* USNM 11320 | Delphinidae | odontocete |
| *Scaphokogia totajpe* MUSM 973 | Kogiidae | odontocete |
| *Schizodelphis barnesi* MNHN AMN 19 | Eurhinodelphinidae | odontocete |
| *Schizodelphis morckhoviensis* USNM 13873 | Eurhinodelphinidae | odontocete |
| *Schizodelphis sp* CCNHM 141 | Eurhinodelphinidae | odontocete |
| *Schizodelphis sulcatus* MGB | Eurhinodelphinidae | odontocete |
| *Semirostrum cerutti* SDNHM 65276 | Phocoenidae | odontocete |
| *Septemtriocetus bosselaersi* IRSNB M.1928 | Phocoenidae | odontocete |
| *Simocetus rayi* USNM 256517 | Simocetidae | odontocete |
| *Sotalia guianensis* USNM 571558 | Delphinidae | odontocete |
| *Sousa chinensis* NHM 1992.97 | Delphinidae | odontocete |
| *Sousa plumbea* USNM 550941 | Delphinidae | odontocete |
| *Sousa sahulensis* NHM 1992.92 | Delphinidae | odontocete |
| *Sousa teuszii* NHM 1992.138 | Delphinidae | odontocete |
| *Squalodon bariensis* IRSNB 2372 | Squalodontidae | odontocete |
| *Squalodon calvertensis* NMNZ MM001996 | Squalodontidae | odontocete |
| *Squalodon* OU 21798 | Squalodontidae | odontocete |
| *Squalodon* OU 22126 | Squalodontidae | odontocete |
| *Squalodon* OU 22397 | Squalodontidae | odontocete |
| *Stenella attenuata* NHM 1966.11.18.5 | Delphinidae | odontocete |
| *Stenella longirostris* USNM 395270 | Delphinidae | odontocete |
| *Steno bredanensis* USNM 572789 | Delphinidae | odontocete |
| Stenodelphininae UCMP 125352 | Lipotidae | odontocete |
| *Tagicetus joneti* IRSNB M. 1892 | Delphinida | odontocete |
| *Tasmacetus shepherdi* USNM 484878 | Ziphiidae | odontocete |
| *Tursiops aduncus* NHM 1882.1.2.3 | Delphinidae | odontocete |
| *Tursiops truncatus gilli* SDNHM 11102 | Delphinidae | odontocete |
| *Tursiops truncatus sp* SDNHM 23798 | Delphinidae | odontocete |
| *Waipatia maerwhenua* OU 22095 | Waipatiidae | odontocete |
| Waipatiid CCNHM 567 | Waipatiidae | odontocete |
| Waipatiid new gen ChM PV7679 | Waipatiidae | odontocete |
| *Xenorophus* new sp ChM PV4823 | Xenorophidae | odontocete |
| *Xenorophus* new sp Yap CCNHM 168 | Xenorophidae | odontocete |
| *Xiphiacetus bossi* USNM 8842 | Eurhinodelphinidae | odontocete |
| *Xiphiacetus cristatus* USNM 21363 | Eurhinodelphinidae | odontocete |
| *Yaquinacetus* USNM 214705 | Squaloziphiidae | odontocete |
| *Zarhachis flagellator* USNM 10911 | Platanistidae | odontocete |
| *Zarhinocetus donnamatsonae* UCMP 86139 | Allodelphinidae | odontocete |
| *Zarhinocetus errabundus* LACM 149588 | Allodelphinidae | odontocete |
| *Ziphius cavirostris* NHM 2006.15 | Ziphiidae | odontocete |

### **Section 1: Additional steps before running geometric morphometric analyses**

#### Missing and variably present bones

We dealt with missing bones in the following way:

#### Missing bones

This refers to bones that should be present but have subsequently broken off or been damaged and could not be reliably digitally reconstructed or mirrored. To estimate positions for landmarks on missing bones, we placed ‘missing’ landmarks as close to the missing structure as possible and then marked it as a ‘missing landmark’ in Checkpoint, which automatically assigns a coordinate of − 9999.

- fixLMtps command from the R package ‘Morpho’ (Schlager, 2017) to estimate missing landmarks by mapping weighted averages from three similar, complete configurations onto the missing specimen.
- Missing landmarks are estimated for missing bones by deforming a sample average or a weighted estimate of the three skull configurations most similar to that with the missing element (Schlager, 2013; Schlager, 2017).
- Estimated landmarks are then added to the deficient configuration (Schlager, 2017). The deformation is performed by a thin-plate-spline interpolation calculated by the available landmarks (Bookstein, 1991; Schlager, 2017).

#### Variably present bones

In *Odobenocetops*, the maxilla does not extend ventrally, and thus must be allocated as ‘absent’ in ventral view. Further, the nasals are absent in Kogiidae (Velez-Juarbe et al., 2015; Benites-Palomino et al., 2019). These absent bones were coded as such for these specimens by placing all relevant landmarks and semi-landmarks onto a single “zero-area” point, adjacent to its position in other taxa, following the method described by Bardua et al. (2019).

#### Resampling and sliding

- We resampled semi-landmark curves to a consistent number with even spacing along each curve across specimens.
- We set semi-landmark numbers to appropriately capture curve shape across the full range of skull shapes, for example to provide suitable sampling of the most dolichocephalic rostra, but to not oversample the most brachycephalic rostra.
- We then slid resampled semi-landmarks along tangents to minimize thin-plate spline bending energy between specimens and the mean shape, resulting in semi-landmark positions that are geometrically homologous across specimens (Gunz et al., 2005; Bardua et al., 2019).
- Following sliding, all morphometric data were subjected to Procrustes superimposition to remove shape variation associated with differences in orientation (both rotation and translation) and isometric size (Rohlf and Slice 1990).

[Table on next page]

### **Table S2.** Description of landmarks placed on each specimen. Includes a description and the number of the corresponding left-hand side (LHS) and right-hand side (RHS) landmarks. Midline landmarks are shown in red.

| **Landmark description** | **Number on LHS of the skull** | **Number on RHS of the skull** |
| --- | --- | --- |
| Nasal anterior | 1 | 67 |
| Left anterior lateral nasal | 2 | 68 |
| Posterior lateral corner of nasal | 3 | 69 |
| Posterior point of nasal | 4 | 70 |
| Tip of rostrum, anterior dorsal side, anterior midline of tooth row (usually premaxilla) | 5 | 71 |
| Anterior dorsal premaxilla | 6 | 72 |
| Posterior dorsal premaxilla | 7 | 73 |
| Anterior lateral ventral premaxilla | 8 | 74 |
| Anterior lateral ventral maxilla | 9 | 75 |
| Dorsal medial maxilla (suture with nasal and premaxilla) | 10 | 76 |
| Nasal-frontal-maxilla suture (posterior medial maxilla) | 11 | 77 |
| Dorsal posterior maxilla on orbit (including lacrimal - dorsal suture frontal) on orbit | 12 | 78 |
| Jugal maxilla, orbit suture - front orbit lateral | 13 | 79 |
| Posterior ventral-lateral most point of maxilla – tooth row - Jugal-maxilla ventral suture | 14 | 80 |
| Posterior tooth row lateral maxilla or lateral maxilla in species with no/negligible dentition | 15 | 81 |
| Jugal anterior dorsal | 16 | 82 |
| Jugal anterior ventral | 17 | 83 |
| Jugal posterior ventral | 18 | 84 |
| Anterior medial frontal | 19 | 85 |
| Posterior medial frontal | 20 | 86 |
| Lateral posterior frontal (posterior lateral parietal suture) | 21 | 87 |
| Postorbital process/bar tip (anterior on crest) | 22 | 88 |
| Anterior lateral frontal (on orbit) | 23 | 89 |
| Anterior dorsal corner of frontal (on orbit) | 24 | 90 |
| Anterior medial parietal | 25 | 91 |
| Posterior medial parietal | 26 | 92 |
| Posterior lateral parietal (squamosal/occipital suture) | 27 | 93 |
| Anterior lateral parietal (on vault) | 28 | 94 |
| Dorsal anterior lateral parietal (suture with frontal) | 29 | 95 |
| Dorsal anterior squamosal suture (with parietal, maybe alisphenoid/frontal) | 30 | 96 |
| Medial anterior zygomatic vault junction (squamosal) | 31 | 97 |
| Anterior dorsal jugal-squamosal suture | 32 | 98 |
| Posterior ventral jugal-squamosal suture, lateral | 33 | 99 |
| Anterior medial most point of the mandibular articular process. | 34 | 100 |
| Posterior lateral most point of the mandibular articular process | 35 | 101 |
| Lateral posterior squamosal (occipital suture) | 36 | 102 |
| Posterior medial dorsal squamosal (parietal/occipital suture) | 37 | 103 |
| MIDLINE: posterior margin of skull roof | *38* | *38* |
| Medial anterior supraoccipital (parietal-occipital suture, usually) | 39 | 104 |
| MIDLINE: dorsal/superior margin of foramen magnum | *40* | *40* |
| Dorsal medial occipital condyle | 41 | 105 |
| Dorsal lateral occipital condyle | 42 | 106 |
| Tip of paraoccipital process - lateral tip | 43 | 107 |
| Lateral ventral occipital + process | 44 | 108 |
| Lateral dorsal occipital | 45 | 109 |
| Ventral medial occipital condyle | 46 | 110 |
| Ventral lateral occipital condyle | 47 | 111 |
| MIDLINE: ventral margin of foramen magnum | *48* | *48* |
| MIDLINE: anterior basioccipital | *49* | *29* |
| Lateral anterior basioccipital | 50 | 112 |
| MIDLINE: anterior most point of basisphenoid, just posterior to the pterygoids and palate | *51* | *52* |
| Lateral anterior basisphenoid | 52 | 113 |
| Lateral posterior basisphenoid | 53 | 114 |
| MIDLINE: Medial posterior basisphenoid | *54* | *54* |
| MIDLINE: Posterior ventral medial point of palate | *55* | *55* |
| MIDLINE: Palatine anterior midline ventral suture | *56* | *56* |
| Pal-pterygoid suture | 57 | 115 |
| Pal-max lateral posterior suture | 58 | 116 |
| Pterygoid posterior | 59 | 117 |
| Ventral posterior pterygoid | 60 | 118 |
| MIDLINE: Maxilla ventral midline posterior suture | *61* | *61* |
| Maxilla ventral midline anterior suture | 62 | 119 |
| Maxilla anterior lateral ventral | 63 | 120 |
| Premaxilla ventral midline posterior suture | 64 | 121 |
| MIDLINE: Anterior-most point of palatal surface immediately posterior to tooth row | 65 | 122 |
| Premaxilla posterior lateral ventral | 66 | 123 |

[Table on next page]

### **Table S3.** Description of landmarks and curves placed on each specimen.

Landmark anchors 1 and 2 are the landmarks between which semi-landmark curves are anchored. I subsampled the curve (‘Resampled curve length’) to ensure all curve points were equally spaced along curves. Midline curves and landmarks are shown in red. The coloured boxes show the following: blue = landmarks and curves placed on the left-hand side (LHS) of the specimens; green = manually placed landmarks on the right-hand side (RHS) of the odontocetes. These (green) landmarks and curves were also placed on the mysticetes but were computer mirrored. Orange = computer mirrored landmarks and curves on mysticetes and odontocetes.

| Semi-landmark curve number | Semi-landmark curve description | Landmark anchor 1 | Landmark anchor 2 | Resampled curve length | Bone |
| --- | --- | --- | --- | --- | --- |
| 1 | Central nasal - anterior to posterior | 1 | 4 | 10 | nasal_l |
| 2 | Posterior nasal – medial to lateral | 4 | 3 | 5 | nasal_l |
| 3 | Lateral nasal – posterior to anterior | 3 | 2 | 10 | nasal_l |
| 4 | Anterior nasal – lateral to medial | 2 | 1 | 5 | nasal_l |
| 5 | Anterior, central, dorsal premaxilla | 5 | 6 | 5 | premax |
| 6 | Anterior to posterior central dorsal premaxilla | 6 | 7 | 35 | premax |
| 7 | Posterior to anterior lateral dorsal premaxilla | 7 | 8 | 35 | premax |
| 8 | Anterior most dorsal maxilla | 8 | 5 | 5 | premax |
| 9 | Medial dorsal maxilla | 9 | 10 | 35 | maxilla |
| 10 | Posterior dorsal maxilla | 10 | 11 | 10 | maxilla |
| 11 | Lateral maxilla over obit | 11 | 12 | 30 | maxilla |
| 12 | Dorsal posterior maxilla on orbit (including lacrimal - dorsal suture frontal) on orbit | 12 | 13 | 10 | maxilla |
| 13 | Jugal maxilla orbit suture - front orbit lateral | 13 | 14 | 10 | maxilla |
| 14 | Posterior ventral lateral most point of maxilla – tooth row - Jugal-maxilla ventral suture | 14 | 15 | 15 | maxilla |
| 15 | Posterior tooth row lateral maxilla or lateral maxilla in species with no/negligible dentition | 15 | 9 | 25 | maxilla |
| 16 | Anterior, medial frontal | 19 | 24 | 30 | frontal |
| 17 | Anterior lateral frontal (on orbit) | 24 | 23 | 10 | frontal |
| 18 | Posterior lateral frontal (on orbit) | 23 | 22 | 10 | frontal |
| 19 | Posterior lateral frontal | 22 | 21 | 15 | frontal |
| 20 | Posterior frontal | 21 | 20 | 20 | frontal |
| 21 | Medial frontal (would be midline in symmetrical taxa) | 20 | 19 | 5 | frontal |
| 22 | Medial dorsal parietal | 25 | 26 | 15 | parietal |
| 23 | Posterior parietal | 26 | 27 | 10 | parietal |
| 24 | Lateral parietal suture with squamosal | 27 | 28 | 15 | parietal |
| 25 | Anterior parietal | 28 | 29 | 15 | parietal |
| 26 | Anterior dorsal parietal | 29 | 25 | 10 | parietal |
| 27 | Anterior squamosal | 32 | 33 | 10 | squamosal |
| 28 | Lateral posterior dorsal squamosal - suture with parietal | 33 | 32 | 25 | squamosal |
| 29 | Medial dorsal zygomatic (with squamosal) – suture with parietal | 31 | 30 | 20 | squamosal |
| 30 | Dorsal posterior zygomatic (with squamosal) | 30 | 37 | 5 | squamosal |
| 31 | Posterior zygomatic (with squamosal) | 37 | 36 | 5 | squamosal |
| 32 | Dorsal medial zygomatic (with squamosal) | 36 | 31 | 20 | squamosal |
| 33 | Anterior ventral mandibular process | 34 | 35 | 15 | mandibular process |
| 34 | Posterior ventral mandibular process | 35 | 34 | 15 | mandibular process |
| *35* | *Midline – Medial supraoccipital* | *39* | *40* | *20* | *supraoccipital* |
| 36 | Supraoccipital suture with dorsal occipital condyle | 40 | 41 | 10 | supraoccipital |
| 37 | Supraoccipital suture with occipital condyle | 41 | 42 | 10 | supraoccipital |
| 38 | Ventral supraoccipital round to exoccipital | 42 | 43 | 15 | supraoccipital |
| 39 | Ventral exoccipital process of supraoccipital | 43 | 44 | 15 | supraoccipital |
| 40 | Dorsal supraoccipital – lateral to medial | 44 | 45 | 20 | supraoccipital |
| 41 | Dorsal medial supraoccipital | 45 | 39 | 10 | supraoccipital |
| 42 | Dorsal medial occipital condyle | 41 | 46 | 15 | occipital condyle |
| 43 | Ventral occipital condyle – medial to lateral | 46 | 47 | 10 | occipital condyle |
| 44 | Lateral occipital condyle | 47 | 41 | 10 | occipital condyle |
| *45* | *Midline – basioccipital medial from posterior to anterior* | *48* | *49* | *20* | *basioccipital* |
| 46 | Anterior basioccipital – medial to lateral | 49 | 50 | 15 | basioccipital |
| 47 | Lateral basioccipital anterior to posterior | 50 | 48 | 25 | basioccipital |
| *48* | *Midline – basisphenoid medial from posterior to anterior* | *54* | *51* | *15* | *basisphenoid* |
| 49 | Anterior basisphenoid medial to lateral | 51 | 52 | 15 | basisphenoid |
| 50 | Lateral basisphenoid – anterior to posterior | 52 | 53 | 15 | basisphenoid |
| 51 | Posterior basisphenoid – lateral to medial | 53 | 54 | 15 | basisphenoid |
| *52* | *Midline – palate posterior to anterior* | *55* | *56* | *20* | *palate* |
| 53 | Anterior palate round to lateral | 56 | 58 | 25 | palate |
| 54 | Posterior palate – lateral to medial | 58 | 57 | 25 | palate |
| 55 | Posterior medial palate – suture with pterygoid | 57 | 55 | 5 | palate |
| 56 | Dorsal posterior pterygoid | 57 | 59 | 10 | pterygoid |
| 57 | Dorsal posterior pterygoid | 59 | 60 | 10 | pterygoid |
| 58 | Anterior pterygoid, medial to lateral | 60 | 57 | 10 | pterygoid |
| 59 | Ventral medial maxilla – posterior to anterior | 61 | 62 | 25 | maxilla_vl |
| 60 | Very anterior lateral of ventral maxilla | 62 | 63 | 5 | maxilla_vl |
| 61 | Lateral maxilla anterior to posterior | 63 | 56 | 45 | maxilla_vl |
| 62 | Lateral ventral premaxilla posterior to anterior | 64 | 66 | 15 | premax |
| 63 | Anterior ventral premaxilla | 66 | 65 | 10 | premax |
| 64 | Medial ventral premaxilla anterior to posterior | 65 | 64 | 15 | premax |
| 65 | Central nasal - anterior to posterior | 67 | 70 | 10 | nasal_r |
| 66 | Posterior nasal – medial to lateral | 70 | 69 | 5 | nasal_r |
| 67 | Lateral nasal – posterior to anterior | 69 | 68 | 10 | nasal_r |
| 68 | Anterior nasal – lateral to medial | 68 | 67 | 5 | nasal_r |
| 69 | Anterior, central, dorsal premaxilla | 71 | 72 | 5 | premax |
| 70 | Anterior to posterior central dorsal premaxilla | 72 | 73 | 35 | premax |
| 71 | Posterior to anterior lateral dorsal premaxilla | 73 | 74 | 35 | premax |
| 72 | Anterior most dorsal maxilla | 74 | 75 | 5 | premax |
| 73 | Medial dorsal maxilla | 75 | 76 | 35 | maxilla |
| 74 | Posterior dorsal maxilla | 76 | 77 | 10 | maxilla |
| 75 | Lateral maxilla over obit | 77 | 78 | 30 | maxilla |
| 76 | Dorsal posterior maxilla on orbit (including lacrimal - dorsal suture frontal) on orbit | 78 | 79 | 10 | maxilla |
| 77 | Jugal maxilla orbit suture - front orbit lateral | 79 | 80 | 10 | maxilla |
| 78 | Posterior ventral-lateral most point of maxilla – tooth row - Jugal-maxilla ventral suture | 80 | 81 | 15 | maxilla |
| 79 | Posterior tooth row lateral maxilla or lateral maxilla in species with no/negligible dentition | 81 | 75 | 25 | maxilla |
| 80 | Anterior, medial frontal | 85 | 90 | 30 | frontal |
| 81 | Anterior lateral frontal (on orbit) | 90 | 89 | 10 | frontal |
| 82 | Posterior lateral frontal (on orbit) | 89 | 88 | 10 | frontal |
| 83 | Posterior lateral frontal | 88 | 87 | 15 | frontal |
| 84 | Posterior frontal | 87 | 86 | 20 | frontal |
| 85 | Medial frontal (would be midline in symmetrical taxa) | 86 | 85 | 5 | frontal |
| 86 | Medial dorsal parietal | 91 | 92 | 15 | parietal |
| 87 | Posterior parietal | 92 | 93 | 10 | parietal |
| 88 | Lateral parietal suture with squamosal | 93 | 94 | 15 | parietal |
| 89 | Anterior parietal | 94 | 95 | 15 | parietal |
| 90 | Anterior dorsal parietal | 95 | 91 | 10 | parietal |
| 91 | Anterior squamosal | 98 | 99 | 25 | squamosal |
| 92 | Lateral posterior dorsal squamosal - suture with parietal | 99 | 98 | 20 | squamosal |
| 93 | Medial dorsal zygomatic (with squamosal) – suture with parietal | 97 | 96 | 5 | squamosal |
| 94 | Dorsal posterior zygomatic (with squamosal) | 96 | 103 | 5 | squamosal |
| 95 | Posterior zygomatic (with squamosal) | 103 | 102 | 20 | squamosal |
| 96 | Dorsal medial zygomatic (with squamosal) | 102 | 97 | 20 | squamosal |
| 97 | Anterior ventral mandibular process | 100 | 101 | 15 | mandibular process |
| 98 | Posterior ventral mandibular process | 101 | 100 | 15 | mandibular process |
| 99 | Supraoccipital suture with dorsal occipital condyle | 40 | 105 | 10 | supraoccipital |
| 100 | Supraoccipital suture with occipital condyle | 105 | 106 | 10 | supraoccipital |
| 101 | Ventral supraoccipital round to exoccipital | 106 | 107 | 15 | supraoccipital |
| 102 | Ventral exoccipital process of supraoccipital | 107 | 108 | 15 | supraoccipital |
| 103 | Dorsal supraoccipital – lateral to medial | 108 | 109 | 20 | supraoccipital |
| 104 | Dorsal medial supraoccipital | 109 | 104 | 10 | supraoccipital |
| 105 | Dorsal medial occipital condyle | 105 | 110 | 15 | occipital condyle |
| 106 | Ventral occipital condyle – medial to lateral | 110 | 111 | 10 | occipital condyle |
| 107 | Lateral occipital condyle | 111 | 105 | 10 | occipital condyle |
| 108 | Anterior basioccipital – medial to lateral | 49 | 112 | 15 | basioccipital |
| 109 | Lateral basioccipital anterior to posterior | 112 | 48 | 25 | basioccipital |
| 110 | Anterior basisphenoid medial to lateral | 51 | 113 | 15 | basisphenoid |
| 111 | Lateral basisphenoid – anterior to posterior | 113 | 114 | 15 | basisphenoid |
| 112 | Posterior basisphenoid – lateral to medial | 114 | 54 | 15 | basisphenoid |
| 113 | Anterior palate round to lateral | 56 | 116 | 25 | palate |
| 114 | Posterior palate – lateral to medial | 116 | 115 | 25 | palate |
| 115 | Posterior medial palate – suture with pterygoid | 115 | 55 | 5 | palate |
| 116 | Dorsal posterior pterygoid | 115 | 117 | 10 | pterygoid |
| 117 | Dorsal posterior pterygoid | 117 | 118 | 10 | pterygoid |
| 118 | Anterior pterygoid, medial to lateral | 118 | 115 | 10 | pterygoid |
| 119 | Ventral medial maxilla – posterior to anterior | 61 | 119 | 25 | maxilla_vl |
| 120 | Very anterior lateral of ventral maxilla | 119 | 120 | 5 | maxilla_vl |
| 121 | Lateral maxilla anterior to posterior | 120 | 56 | 45 | maxilla_vl |
| 122 | Lateral ventral premaxilla posterior to anterior | 121 | 123 | 15 | premax |
| 123 | Anterior ventral premaxilla | 123 | 122 | 10 | premax |
| 124 | Medial ventral premaxilla anterior to posterior | 122 | 121 | 15 | premax |
